# Stiffness and Viscoelasticity of Adipose Tissue Decellularized Extracellular Matrix Hydrogels Influence Proliferation, Growth Pattern, Migration and Invasion of Breast Cancer Cells

**DOI:** 10.64898/2025.12.11.693682

**Authors:** Naike Salvador Moreno, Maria Mercedes Fernandez, Mikel Azkargorta, Iratxe Madarieta, Roberto Hernandez, Felix Elortza, Beatriz Olalde, Amaia Cipitria

**Affiliations:** Group of Bioengineering in Regeneration and Cancer, Biogipuzkoa Health Research Institute, San Sebastian, Spain.; POLYMAT Institute, University of the Basque Country -EHU, San Sebastián, Spain; Proteomics Platform, CIC BioGUNE, Basque Research and Technology Alliance (BRTA), Derio, Spain; TECNALIA, Basque Research and Technology Alliance (BRTA), San Sebastian, Spain; Group of Bioengineering in Regeneration and Cancer, Biogipuzkoa Health Research Institute, San Sebastian, Spain; IKERBASQUE, Basque Foundation for Science, Bilbao, Spain.

**Author notes:** Corresponding author: N. Salvador Moreno,; A. Cipitria.

**Keywords:** Decellularized extracellular matrices (dECMs), elasticity, viscoelasticity, dECM proteomic analysis, breast cancer, growth patterns, invasion

## Abstract

Breast cancer remains the leading cause of cancer-related mortality among women. Cellular behavior is influenced by the physicochemical factors of the extracellular matrix (ECM) surrounding them. In order to model the breast ECM, adipose tissue decellularized extracellular matrix (dECM)-derived hydrogels are generated, which retain the full biochemical complexity of the source tissue, while also exhibiting stiffness and viscoelastic properties comparable to those of breast tissue. By recapitulating the characteristics of their native environment, validated through proteomic analysis and rheology, the poorly metastatic MCF-7 and the highly invasive MDA-MB-231 breast cancer cell lines exhibit their archetypal behaviors within the 3D hydrogels. Moreover, these cells display significantly distinct proliferation, invasion, and growth patterns in hydrogels with different stiffness and viscoelasticity, highlighting the importance of biophysical parameters in modulating cell phenotype. These results illustrate that user-friendly 3D biomaterial models based on adipose tissue dECM can effectively replicate crucial aspects of in vivo cellular behavior.

**Table of Contents:** This study presents 3D hydrogels derived from porcine adipose tissue dECM that mimic biophysical and biochemical properties of breast tissue. Two breast cancer cell lines mimicking luminal and triple-negative subtypes, exhibited distinct proliferation, migration, and invasion behaviors depending on hydrogel stiffness and viscoelasticity. The results highlight how biophysical cues shape cell phenotype in a user-friendly 3D model.

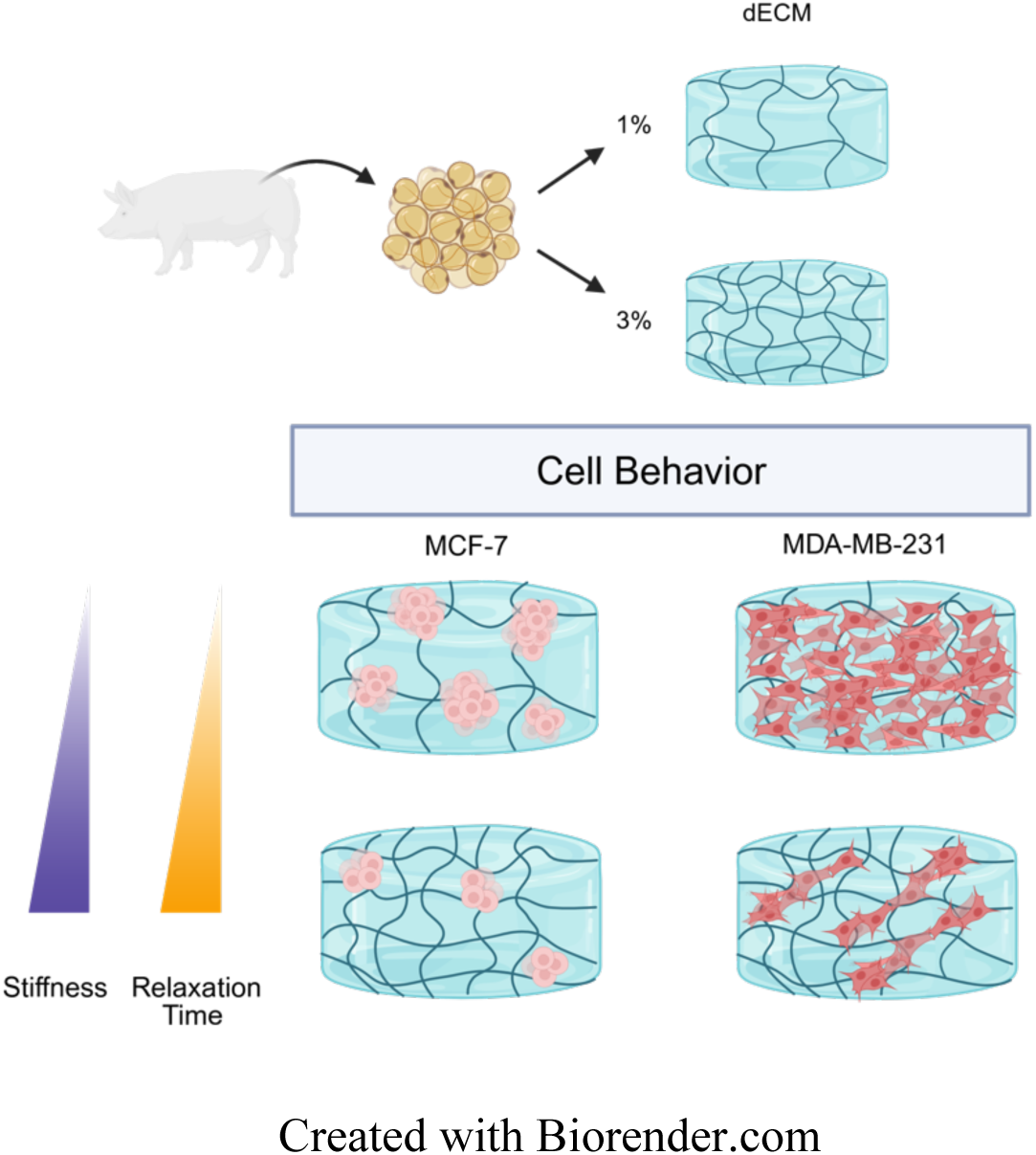

Created with Biorender.com

## 1. Introduction

Creating an environment that accurately mimics the native breast extracellular matrix (ECM) is essential for developing effective in vitro breast tumor models. In the human body, the breast is among the softest tissues. The storage modulus of normal breast tissue has been reported to range between 160 Pa to 950 Pa.^[1–3]^ This variability arises from differences between individuals, variations between different breast compartments (fatty and fibroglandular tissue), and the diverse methodologies used to measure tissue stiffness, such as nanoindentation or various types of elastography (including ultrasound elastography or magnetic resonance elastography (MRE) which utilizes MRI). ^[4]^

Cellular behavior is profoundly modulated by the mechanical properties of the surrounding ECM, primarily through mechanotransduction signaling pathways. The cytoskeleton transmits mechanical signals to the cell nucleus, ^[5]^ rapidly converting these cues into biochemical signals and metabolic responses. ^[2]^ On compliant materials, cells deform the substrate, while on stiffer substrates, the cells themselves deform, which modifies cell cycle, proliferation and migration. ^[6]^ During pathogenesis and tumor progression, alterations in mechanical properties, ^[7,8]^ such as increased stiffness, occur. These changes are primarily due to a higher amount of crosslinked and oriented collagen. ^[9]^ For instance, in breast cancer, the tissue’s elastic modulus can more than double as the tumor progresses. ^[10]^

Three-dimensional (3D) tumor models for breast cancer, which replicate key physical and biochemical aspects of the tumor microenvironment, are promising tools for studying cell behavior in a physiologically relevant context and for developing personalized therapeutic applications. ^[11,12]^ Unlike traditional two-dimensional (2D) cell cultures, 3D organotypic systems can mimic specific biochemical and biophysical aspects, such as stiffness or viscoelasticity, allowing for precise customization to address various research questions. Most studies employ hydrogels with a storage modulus ranging from just over 4 kPa to 20 kPa to assess the proliferation and migration capacity of cancer cells. ^[13]^ This range, while informative, diverges considerably from the mechanical properties of native breast tissue.

Although research has primarily focused on stiffness, recognizing it as a hallmark of many cancer types, tissues exhibit complex mechanical behaviors. Therefore, it is important to study other aspects, such as viscoelasticity, often characterized by stress relaxation assays. Viscoelastic materials, which display both liquid-like (viscous) and solid-like (elastic) properties, respond to mechanical stimuli with an initial elastic reaction followed by a time-dependent dissipation of energy. ^[14]^ Research has highlighted the importance of matrix stress relaxation as a fundamental property of physiological ECMs, crucial for understanding cell-ECM interactions and the biophysics of mechanotransduction that influences cell spreading. ^[15,16]^ Despite using different dimensionality and cell types, it has been found that cell morphology and spreading vary significantly on substrates exhibiting stress relaxation compared to purely elastic substrates. ^[17,18]^ Increasing evidence suggests that changes in tissue viscoelasticity may correlate with cancer progression. ^[19,20]^ In addition, recent studies suggest that tissue viscoelasticity could serve as a cancer biomarker, with tissue fluidity facilitating the differentiation between normal, benign, and malignant tumors. ^[21,22]^ Nevertheless, it is important to be aware that stress relaxation times are not absolute measures due to the absence of standardized protocols for the mechanical characterization of tissues and hydrogels. In the few studies that measure the viscoelastic properties of tumors, the most frequently used techniques are based on nanoindentation, ^[4,23]^ with oscillatory rheometry and compression tests being exceptions. However, these latter techniques are more commonly applied for hydrogel characterization.

The tissue ECM architecture and composition have proven to be as important. Specifically, collagen alignment and density have been correlated with breast tumor aggressiveness and the invasiveness of carcinoma cells, while properties like collagen fibril diameter additionally regulate cell cluster formation and invasion. ^[24,25]^ Prior research has also shown that cell transmigration across collagen interfaces, characterized by changes in collagen directionality, pore size, or both, induce a more aggressive phenotype and alters the migration path. ^[26]^

Single-component scaffolds (such as gelatin, collagen, or alginate) are common models to study biomechanical phenomena, as they can accurately recapitulate specific properties of tissues, such as stiffness. ^[27]^ However, they poorly mimic the complexity of native breast tissue. In this sense, adipose tissue decellularized extracellular matrices (dECMs) can better replicate the biochemical and biophysical characteristics of native ECM. They do so without incorporating immunostimulatory nucleic acids and tumor-stimulatory factors, which are commonly found in tumor-derived ECMs like Matrigel. ^[28]^

In order to better understand the dynamic interplay between different cancer cells and their respective microenvironment during the initial stages of tumor growth and invasion, we generated a biomimetic ECM model that recapitulates the biomechanics and biochemistry of breast tissue. For this purpose, we chose porcine adipose tissue due to its abundance, similarity to human breast tissue in terms of ECM composition, and biocompatibility with human-derived cells. ^[29,30]^ By studying two representative matrices in terms of stiffness and viscoelasticity, and sequentially combining them, we were able to observe the phenotype transitions of the poorly metastatic MCF-7 and the highly invasive MDA-MB-231 breast cancer cell lines. These tissue inspired models served as an effective platform to investigate the link between ECM properties and breast cancer cell behavior.

In this study, we aimed to model the mechanical properties and structural protein composition of the breast microenvironment to explore in more detail the possible triggers of behavioral changes of breast cancer cells, such as variations in proliferation and migration patterns. In addition, our investigation of the proteins present in the source dECM powder, compared with those in the dECM-derived hydrogels, revealed that the proteins encountered by the cells within the dECM hydrogels are predominantly associated with structural support and cell adhesion functions. Using artificial intelligence models to analyze our results, we provide functional evidence of proliferation and cell cycle phase changes in MDA-MB-231 and MCF-7 cells growing embedded in hydrogels that exhibit distinct viscoelastic properties, which matches previously described phenotypes. Moreover, we gained deeper insight into how variations in both stiffness and viscoelasticity influence the growth pattern of different cell types. These results enhance our understanding of the biophysical and biochemical factors that drive proliferative and invasive phenotypes of breast cancer cells, underscoring the critical role of ECM biomechanical properties in this process.

## 2. Results

### 2.1. Adipose tissue dECM-derived hydrogels preserve a significant degree of structural, adhesion and secreted proteins from the source tissue

Traditional 2D cultures lack the biochemical, physical and structural cues of the native tissue ECM. To overcome these limitations, 3D matrices and, in particular, dECMs are gaining attention because they can partially retain the biochemical and structural properties of the source tissue. dECMs possess a complex biochemical composition, containing proteins essential for ECM assembly and influencing key cellular processes such as adhesion, growth, differentiation, and signal transduction. To identify the proteins that interact with cells growing within the dECM-derived hydrogels, we conducted a liquid chromatography– tandem mass spectrometry (LC-MS/MS) analysis. We analyzed and compared the protein content of the source adipose tissue dECM powder obtained from Tecnalia with that of the adipose tissue dECM-derived hydrogels, which allowed us to evaluate how processing into a hydrogel alters the protein content of the adipose tissue dECM (**Figure 1**).

**Figure 1.**
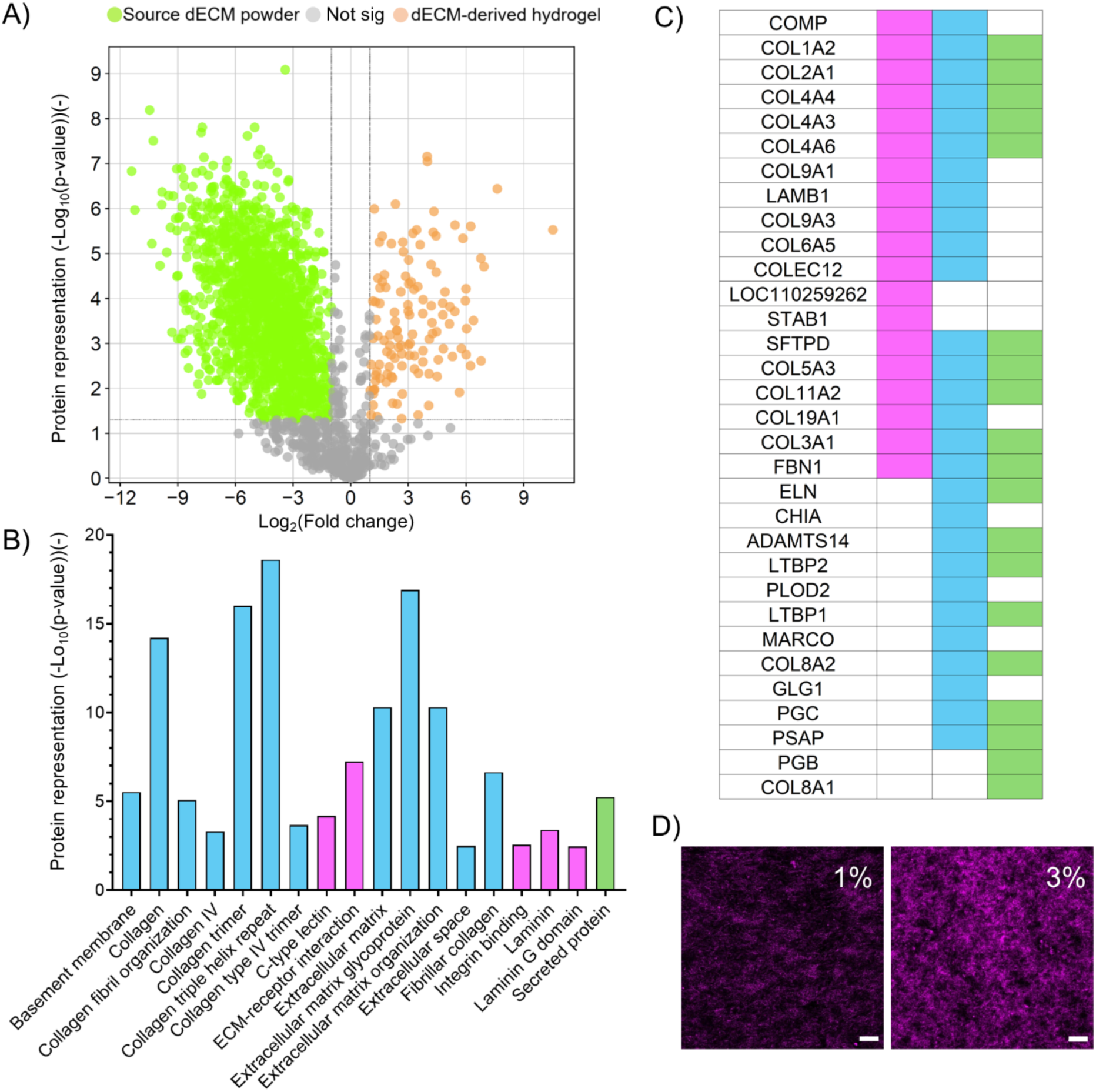
A) Volcano plot representing proteins present in the adipose tissue dECM-derived hydrogel (orange dots) compared to the source dECM powder (green dots) using log2 (fold change) as the x-axis and protein representation (−log10 (p-value)) as the y-axis. Not significant dots are depicted in grey. B) Functional enrichment of the proteins significantly represented in the adipose tissue dECM-derived hydrogels using the protein representation (-log10 (p-value)) as the y-axis. C) Table of the genes coding for proteins significantly represented in the adipose tissue dECM-derived hydrogels grouped into three categories: cellular adhesion (pink), structural ECM (blue), and secreted proteins (green). D) Representative confocal reflectance images of the collagen fibers (purple) in the 1% and 3% adipose tissue dECM-derived hydrogels. For both 1% and 3% hydrogels, n=3. The excitation and emission wavelengths were 633 and 620-640 nm. Scale bar: 20 μm.

As shown in the volcano plot (Figure 1A), a significant number of proteins were more abundant in the source dECM powder but showed reduced detectability in the dECM-derived hydrogel. This finding supports previous studies ^[31]^ that reported a 50% reduction in the total number of proteins in adipose tissue dECM-derived hydrogels due to processing. The major loss is attributed to the acidic digestion of the dECM powder with pepsin, which cleaves phenylalanine-phenylalanine and phenylalanine-tyrosine bonds.

Given that cells proliferate within dECM-derived hydrogels and are influenced by their protein composition, we focused on characterizing proteins that were significantly more abundant in this material. The genes encoding these proteins were functionally annotated using the DAVID program. The protein representation, or in other words, the significance of each function, is visualized in the bar graph (Figure 1B). Because many functions were closely related, we grouped similar functions into three categories: cellular adhesion (pink), structural ECM (blue), and secreted proteins (green). The table in Figure 1C shows which genes are associated with each function. Of these, 42% had structural functions (blue), 29% were involved in cellular adhesion (pink), and 29% were secreted proteins (green). Most of the identified proteins were associated with collagen formation, trimerization, and ECM glycoproteins. This is coherent since collagen type I is the primary component of organized fiber structures in most connective tissues. ^[32]^ It is a heterotrimer comprised of two collagen alpha-1 chain units and one unit of collagen alpha-2, encoded by the COL1A1 and COL1A2 genes, respectively. ^[33]^ Collagen is resistant to most protease activity due to the primary sequence of the α chains, which are rich in Gly, Pro, and Hyp, and also because of its triple-helix structure, which conceals cleavage sites in cryptic pockets. ^[34]^ Thus, after decellularization most collagen content has been shown to be preserved, which contributes to the adhesion of cells and other ECM components. ^[35]^ To analyze the structure of the collagen in the adipose tissue dECM-derived hydrogel, we utilized confocal reflectance microscopy. With this technique, the wavelength used to illuminate the sample is the same as the one received from the sample. This reveals the orientation of the collagen fibers forming the fibrillar architecture of the dECM-derived hydrogels. ^[36]^ We observe, as expected, a higher density of fibers in the 3% adipose tissue dECM hydrogel (Figure 1D). However, there does not appear to be a preferential direction of fiber orientation within the adipose tissue dECM-derived hydrogels. Proteins related to lectin and laminin also play a primary role in cellular adhesion, allowing cell attachment and migration. Additionally, many secreted proteins, such as fibrillin, are linked to structural and adhesion functions as well. Altogether, the presence of these molecules and components in the adipose tissue dECM-derived hydrogels, which retain the biochemical properties of the source tissue to a great extent, supports their use as a biologically relevant ECM.

### 2.2. Adipose tissue dECM-derived hydrogels exhibit stiffness and stress relaxation times in a range comparable to those of native breast tissue

In addition to retaining biochemical composition of the native adipose tissue, we also aimed to replicate its biophysical properties and thereby create a biomimetic 3D environment. Mimicking their natural microenvironment ensures that cells behave as closely as possible to their natural state within the human body. The breast is one of the softest tissues in the human body, with an elastic modulus E less than 1 kPa.^[1]^ Consequently, we generated adipose tissue dECM-derived hydrogels at two different concentrations, 1% and 3% w/v, to approximate stiffness and stress-relaxation times of native breast tissue.

We conducted frequency sweep tests within the linear viscoelastic region for 1% and 3% adipose tissue dECM-derived hydrogels, which revealed complex moduli *G\** of 60 Pa and 300 Pa, corresponding to elastic moduli *E* of 180 Pa and 900 Pa, respectively (**Figure 2A** and **B**). This demonstrates that, despite their statistically significant differences in stiffness, both hydrogels still replicate the mechanical characteristics of the natural breast microenvironment. The phase angle tangent was very similar for both gels, approximately 0.2, indicating a predominantly elastic nature (tan(δ)=0) rather than viscous (tan(δ)=1) (Figure 2C). Hydrogels containing cells exhibited comparable values (Figure S1C), reflecting closely matched mechanical behavior.

**Figure 2.**
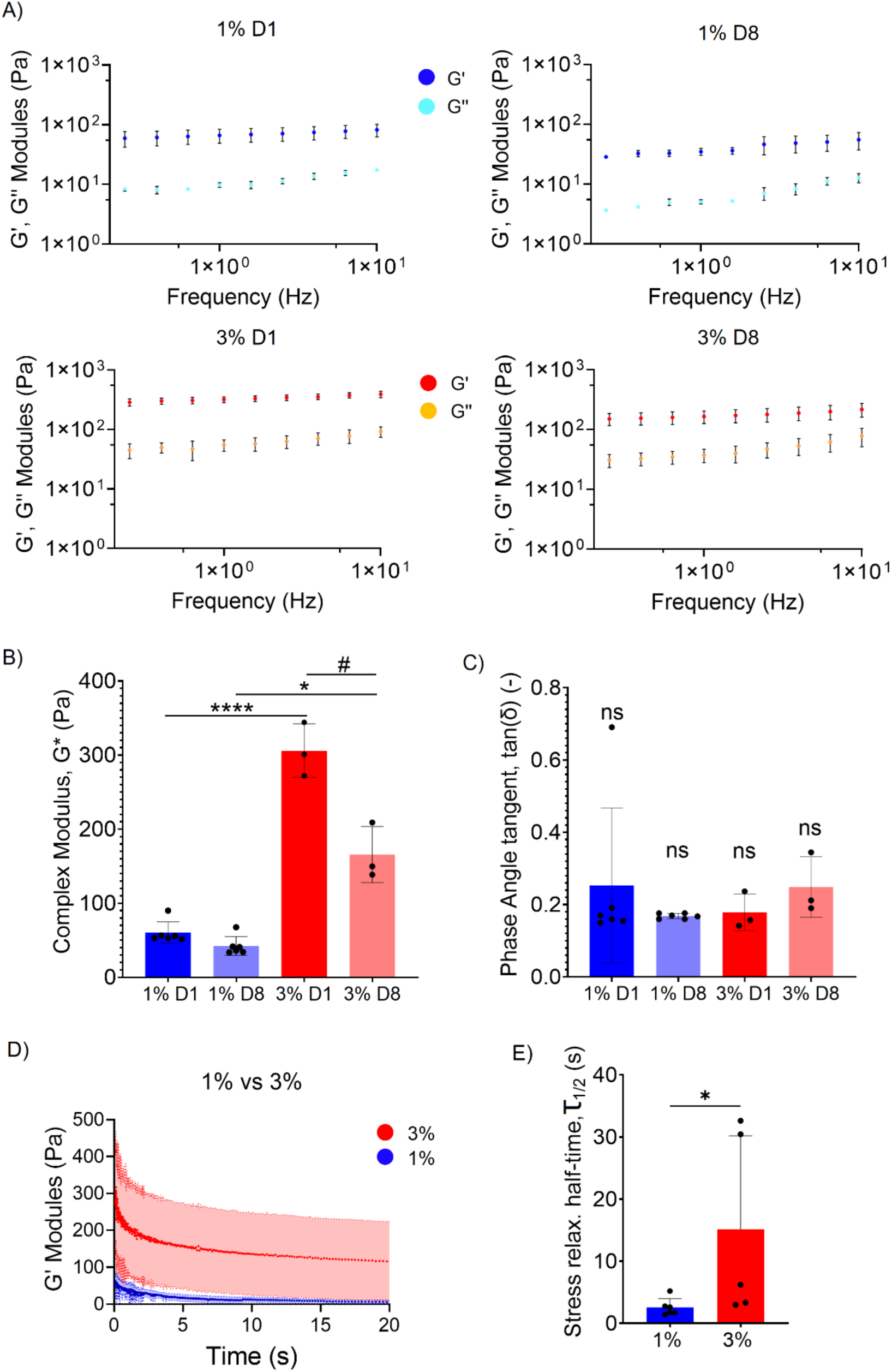
A) Frequency sweep tests (from 0.02 to 10 Hz) of the 1% and 3% adipose tissue dECM-derived hydrogels performed at constant strain (0.5%) at day 1 and 8. n=6 for 1% hydrogels and n=3 for 3% hydrogels. Storage (*G′*) and loss (*G″*) moduli are represented as colored and fainter circles, respectively. B) Complex modulus (*G\**) and C) phase angle tangent (tan(δ)) of 1% and 3% dECM-derived hydrogels at day 1 and 8. Data were calculated from the frequency sweep tests in A. D) Stress relaxation tests (at 2% shear strain) and E) stress relaxation half-times (*τ*_1/2_) of the 1% and 3% adipose tissue dECM-derived hydrogels at day 1. n=5 for 1% hydrogels and n=5 for 3% hydrogels. Data were analyzed using an unpaired Student’s t-test: * (p< 0.05).

To investigate whether the hydrogels underwent degradation over time, we performed frequency sweeps on day 1 and 8 post-synthesis (Figure 2A). Although the 1% gels showed minimal differences in stiffness between days 1 and 8, the 3% gels exhibited some intrinsic degradation, as their complex modulus significantly decreased by approximately 100 Pa (Figures 2A and 2B). We also examined whether the cellular activity within the gels would accelerate the degradation of the adipose tissue dECM-derived hydrogels over time. However, no significant differences were observed in the frequency sweeps of 1% hydrogels with and without cells conducted on day 1 and day 8 (Figure S1A and B).

To further characterize the viscoelastic behavior of the adipose tissue dECM-derived hydrogels we performed stress-relaxation tests. The 1% hydrogels exhibited a significantly faster relaxation compared to the 3% hydrogels (Figure 2D and E), with mean values of stress relaxation half-times (*τ*_1/2_) of 2.3 and 14.8 seconds, respectively (Figure 2E and Figure S1D). This reveals that, together with stiffness, the relatively small viscous component is primarily responsible for the differences in the mechanical properties of the hydrogels.

### 2.3. Breast cancer cells exhibited distinct proliferation, growth patterns, migration and invasion capacity in 1% vs 3% adipose tissue dECM-derived hydrogels

To investigate cellular behavior in these dECM-derived hydrogels that mimic the mechanical and biochemical properties of breast tissue, we analyzed three key aspects: cell proliferation, growth patterns, migration and invasion.

#### 2.3.1. Breast cancer cells proliferate more in the softest and fastest relaxing 1% adipose tissue dECM-derived hydrogels

To study cell proliferation, we selected two well-characterized breast cancers cell lines with differing aggressiveness: MCF-7 and MDA-MB-231. We used genetically engineered cells expressing FUCCI2, which allows real-time quantification and visualization of cell cycle progression, distinguishing between the G1 phase (red), and the G2/M phase (green), without additional staining (**Figure 3B** and **4B**).

**Figure 3.**
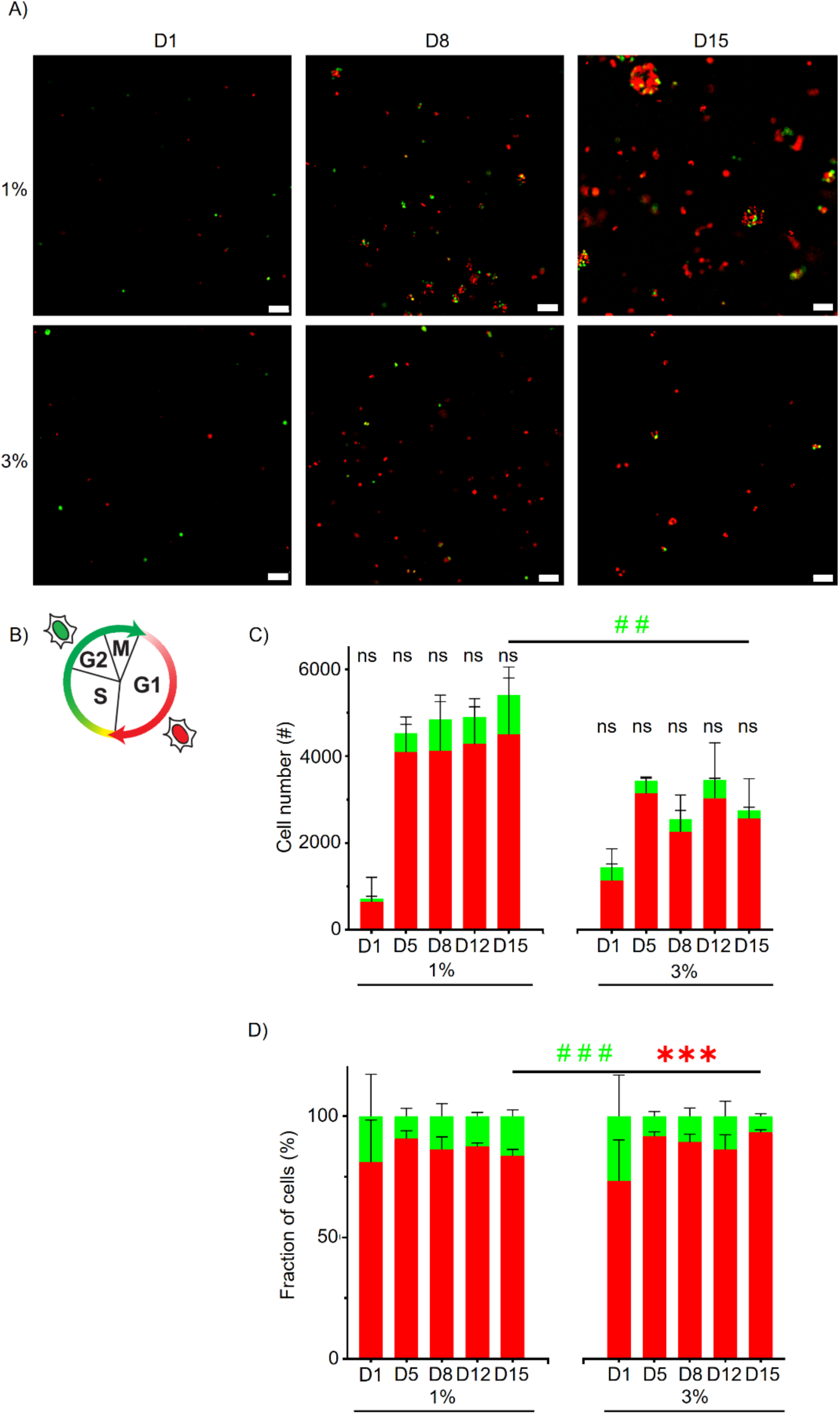
A) Fluorescence microscope images of MCF-7 cells expressing FUCCI2 (G1 shown in red, G2/M shown in green) growing within 1% and 3% adipose tissue dECM-derived hydrogels after 1, 8 and 15 days in culture. Scale bar: 100 μm. B) Schematic representation of the color of the FUCCI-modified cell nucleus progressing through the cell cycle (G1: red, G2/M: green) adapted from ^[37]^, 2008, with permission from Elsevier. C) Quantification of the number of cells in the G1 phase or the G2/M phase at days 1, 5, 8, 12 and 15 in 1% and 3% adipose tissue dECM-derived hydrogels using Dragonfly. D) Fraction of cells in either the G1 phase or the G2/M phase for each time point and material. Experiments were repeated three times per condition. Data were analyzed using an unpaired Student’s t-test: n.s (p > 0.05), ** (p < 0.01), *** (p< 0.001).

**Figure 4.**
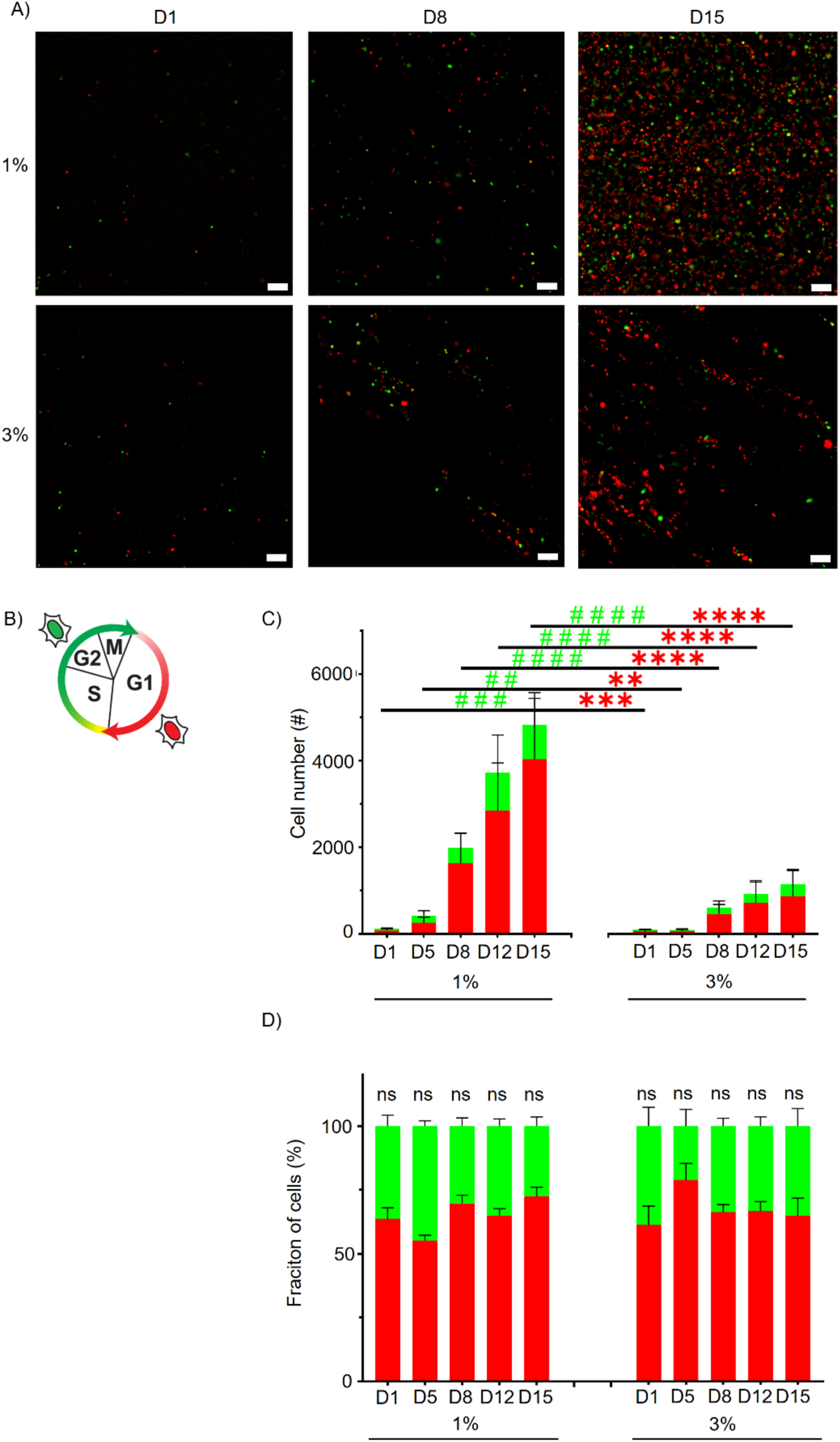
A) Fluorescence microscope images of MDA-MB-231 cells expressing FUCCI2 (G1 shown in red, G2/M shown in green) growing within 1% and 3% adipose tissue dECM-derived hydrogels after 1, 8 and 15 in culture. Scale bar: 100 μm. B) Schematic representation of the color of the FUCCI-modified cell nucleus progressing through the cell cycle (G1, red; G2/M, green) adapted from **^[^**^37]^, 2008, with permission from Elsevier. C) Quantification of the number of cells in the G1 phase or the G2/M phase at days 1, 5, 8, 12 and 15 in 1% and 3% adipose tissue dECM-derived hydrogels using Dragonfly software. D) Fraction of cells in either the G1 phase or the G2/M phase for each time point and material. Experiments were repeated three times per condition. Data were analyzed using an unpaired Student’s t-test: n.s (p > 0.05), ** (p < 0.01), *** (p< 0.001), **** (p< 0.0001).

MCF-7 and MDA-MB-231 cells were cultured within 1% and 3% adipose tissue dECM-derived hydrogels over a period of 15 days, with images captured at 1, 5, 8, 12, and 15 days. It is noteworthy that these hydrogels gellify thermally, which avoids UV-induced DNA damage. Using Dragonfly software, we trained a model to identify and quantify cells in the G1 and G2/M phases. This enabled us to calculate both cell proliferation rates and the proportion of cells in each phase of the cell cycle for MCF-7 (Figure 3C and D) and MDA-MB-231 (**Figure 4C** and **D**).

Both MCF-7 and MDA-MB-231 cells displayed representative behavior of luminal and triple-negative breast cancer, respectively; MCF-7 cells initially grew at a noticeable rate but then slowed down and the number of cells stabilized (Figure 3A and C), while MDA-MB-231 cells presented exponential growth, with a rapid increase in cell number that was sustained throughout the experiment (Figure 4A and C).

Interestingly, even though the complex moduli of the adipose tissue dECM-derived hydrogels were within the range considered ‘soft’ (*G\** of 60 Pa for 1% and 300 Pa for 3%) and viscoelastic (*τ*_1/2_ of 2.3 and 14.8 seconds for 1% and 3% hydrogels, respectively), we observed striking differences in the behavior of MCF-7 and MDA-MB-231 cells cultured in 1% vs 3% hydrogels across all tested parameters. MCF-7 cells exhibited enhanced proliferation at day 15 in the softest and fastest-relaxing 1% adipose tissue dECM-derived hydrogels compared to the 3% hydrogels (Figure 3C), and for MDA-MB-231, such differences were significant at all timepoints (Figure 4C).

Regarding cell cycle distribution, MCF-7 cells showed significant differences in the fraction of cells in the G1 phase (red cells) versus the G2/M phase (green cells) at later timepoints when comparing 1% and 3% adipose tissue dECM-derived hydrogels (Figure 3D). The 3% hydrogels displayed a higher G1-to-G2/M ratio, indicating that more MCF-7 cells were in the G1 phase, suggesting a trend toward growth arrest. In contrast, MDA-MB-231 cells displayed no significant differences in cell cycle phase distribution across materials and time points, confirming they retained their highly proliferative phenotype (Figure 4D).

#### 2.3.2. Breast cancer cells exhibit significantly distinct growth patterns in 1% vs 3% adipose tissue dECM-derived hydrogels

Given the observed differences in proliferation between MCF-7 and MDA-MB-231 cells grown within 1% and 3% adipose tissue dECM-derived hydrogels, we proceeded to delve into the study of breast cancer cell growth pattern.

MCF-7 cells were cultured within 1% and 3% adipose tissue dECM-derived hydrogels over a period of 15 days. Using Dragonfly software, we trained a model to identify and measure spheroids. Since the average epithelial cell is 15 µm long, we established a lower limit of 30 µm for spheroid size, assuming that a spheroid must consist of at least two cells (Figure S2A and B). After 15 days in culture, there were remarkably more spheroids and of bigger size in the 1% compared to the 3% adipose tissue dECM-derived hydrogels (**Figure 5A** and **B**).

**Figure 5.**
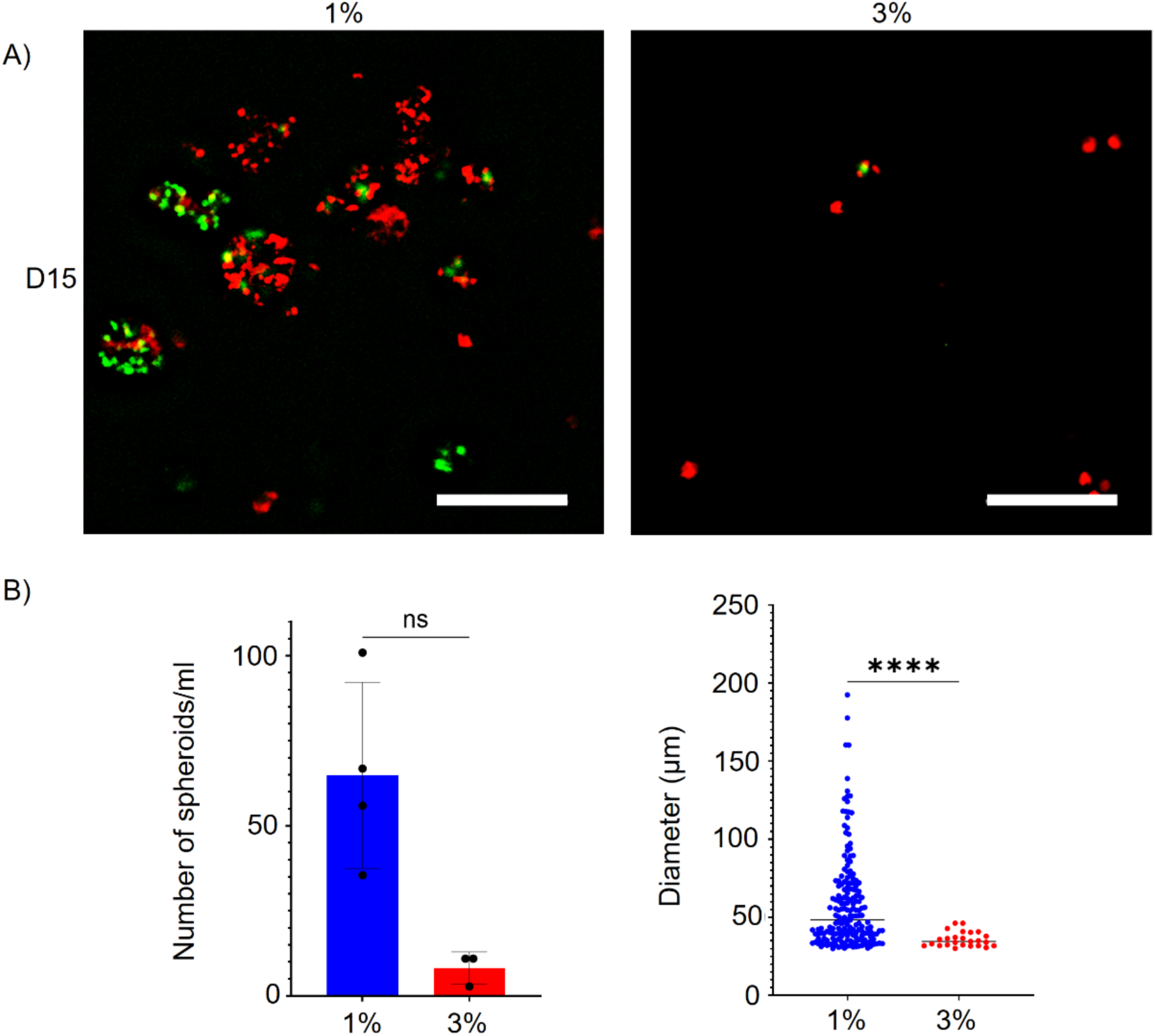
A) Fluorescent microscope images of MCF-7 cells expressing FUCCI2 (G1: red, G2/M: green) growing within 1% and 3% adipose tissue dECM-derived hydrogels after 15 days in culture. Scale bar: 100 μm. B) Quantification of the number and size of spheroids in maximum projection images of 100 μm-thick z-stacks from four and three replicates for 1% (blue) and 3% (red) hydrogels, respectively, after 15 days in culture, using Dragonfly software. Data were analyzed using an unpaired Student’s t-test: * (p < 0.05), **** (p< 0.0001).

MDA-MB-231 cells were cultured for 15 days within 1% and 3% adipose tissue dECM-derived hydrogels. In this case, to be able to see the structure of the cytoskeleton and the nucleus, the parental cells were stained with DAPI and phalloidin on days 1, 8 and 15. The cells showed markedly different growth patterns in 1% and 3% hydrogels (Figure 6A and Figure S3A and B). This time Dragonfly software was used to train an artificial intelligence model to define the growth direction of a connected group of cells by calculating the angle between the X-axis and the group’s major axis (Figure S3C). The histograms (**Figure 6B**) display the number of cell clusters oriented towards specific angles. In the 3% adipose tissue dECM-derived hydrogels, there is a clear tendency for cells to grow in a preferential direction. In contrast, the 1% hydrogels show no preferred growth direction for the cells.

**Figure 6.**
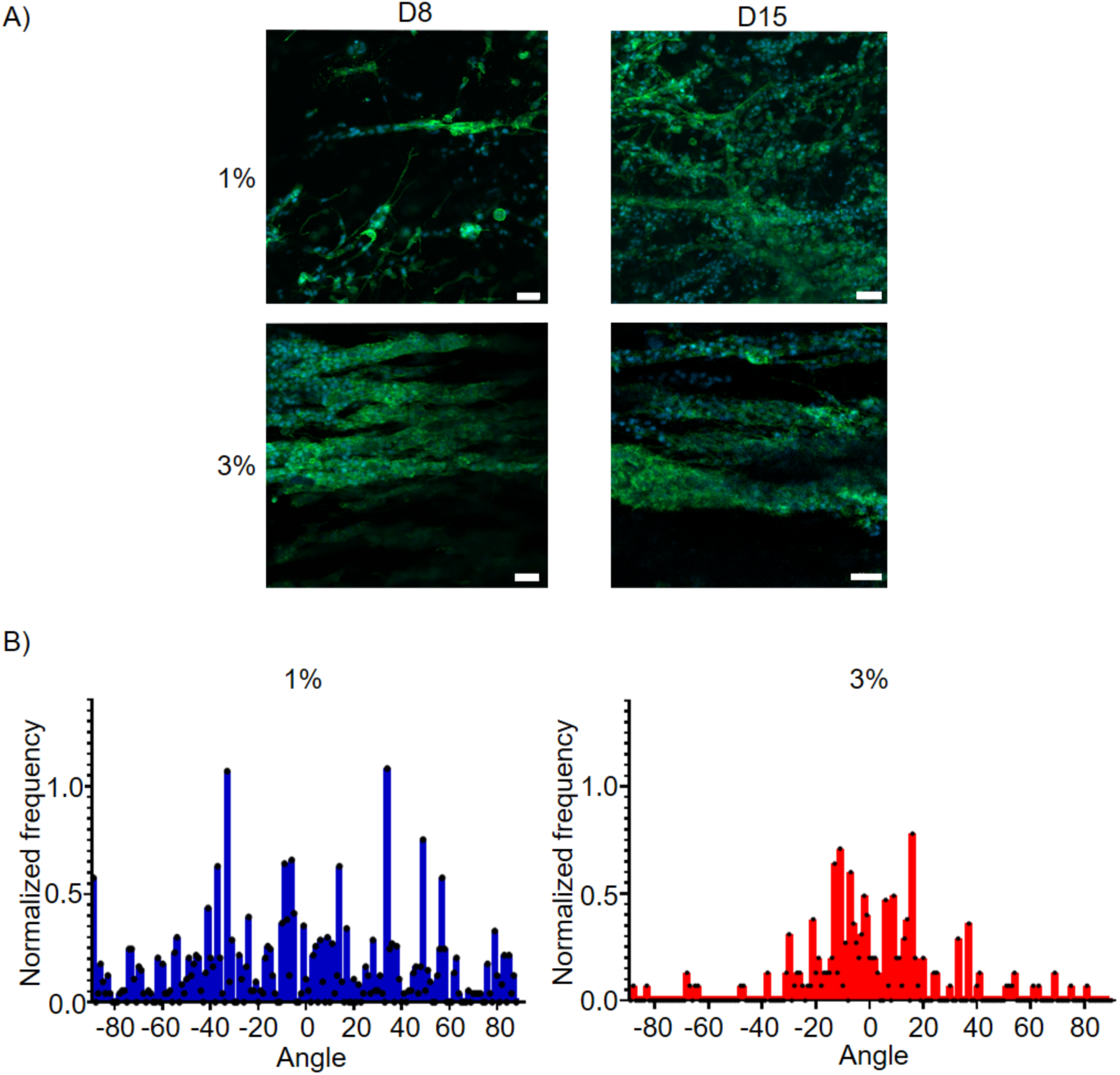
A) Confocal microscope images of MDA-MB-231 cells stained with DAPI (blue) and phalloidin (green) growing within 1% and 3% adipose tissue dECM-derived hydrogels after 8 and 15 days in culture. Experiments were repeated six times per condition. Scale bar: 50 μm. B) Histograms showing the growth angles of connected cells in 1% (blue) and 3% (red) adipose tissue dECM-derived hydrogels after 15 days in culture, quantified using Dragonfly software.

#### 2.3.3. Higher number of MDA-MB-231 cells migrate and invade the softest and fastest relaxing 1% adipose tissue dECM-derived hydrogels

Given the differences observed in cell proliferation and growth pattern between 1% and 3% adipose tissue dECM-derived hydrogels, we further assessed the migration and invasion capacity of MCF-7 and MDA-MB-231 cells employing the commercial microfluidic system OrganoPlate® 3-lane 40 (Mimetas®). The inclusion of a simple perfusion system enhances its physiological relevance. (Figure S4A and B). To study the capacity of cells to migrate and invade from one hydrogel to another, cells were cultured in microfluidic chips containing cell-laden 1% and 3% adipose tissue dECM-derived hydrogels placed in direct contact with cell-free hydrogels of the opposite concentration. Specifically, 1% hydrogels containing cells were paired with cell-free 3% hydrogels, and vice versa (**Figure 7A** and **8A**). The interface boundary was defined at the time of polymerization, as the margin between hydrogels was visible based on phase contrast (Figures 7B, C, 8C and D). The distance from the interface boundary to structural components of the chip (the tip of the triangle on top of the central channel, grey trapezoid in Figures 7B, C and 8C, and white rectangles in Figure 8D) was measured and used as a reference in subsequent timepoints. Chips with homogeneous adipose tissue dECM-derived hydrogels seeded with cells were kept as growth controls (Figure S5A and B). Images of the invasion process were captured at different time points (Figures 7B, C, 8C and D), and the distance invaded by individual cells from the original interface into the acellular hydrogel was quantified at day 8 using Fiji (Figure 8B).

**Figure 7.**
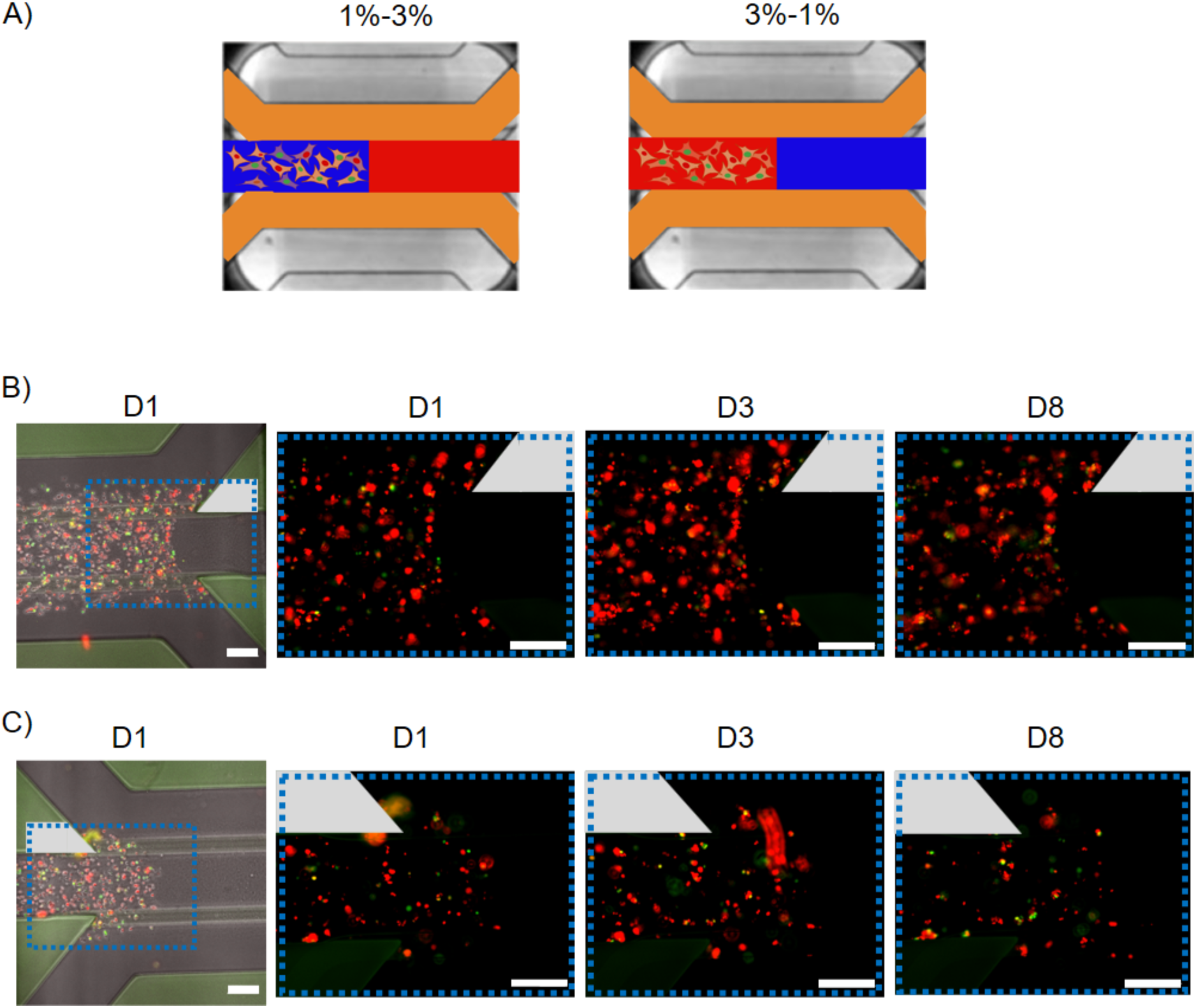
A) Schematic representation of the interphase between 1% (blue) adipose tissue dECM-derived hydrogels embedded with cells and 3% (red) cell-free adipose tissue dECM-derived hydrogels, and vice versa, 3% adipose tissue dECM hydrogels with cells and cell-free 1%, in the central channel. Bright field and fluorescence microscope images of MCF-7 cells expressing FUCCI2 (G1: red, G2/M: green) in the area of interest, B) including regions from the 1% and the 3% adipose tissue dECM-derived hydrogels and their interphase (1%-3%) after 1, 3 and 8 days, and C) including regions from the 3% and the 1% adipose tissue dECM-derived hydrogels and their interphase (3%-1%) after 1, 3 and 8 days. The grey trapezoid highlights the structural guide used as a reference for cell migration. The blue dashed line indicates the zoom-out area imaged at higher magnification captured at subsequent timepoints. Experiments were repeated two times per condition. Scale bar: 200 μm.

**Figure 8.**
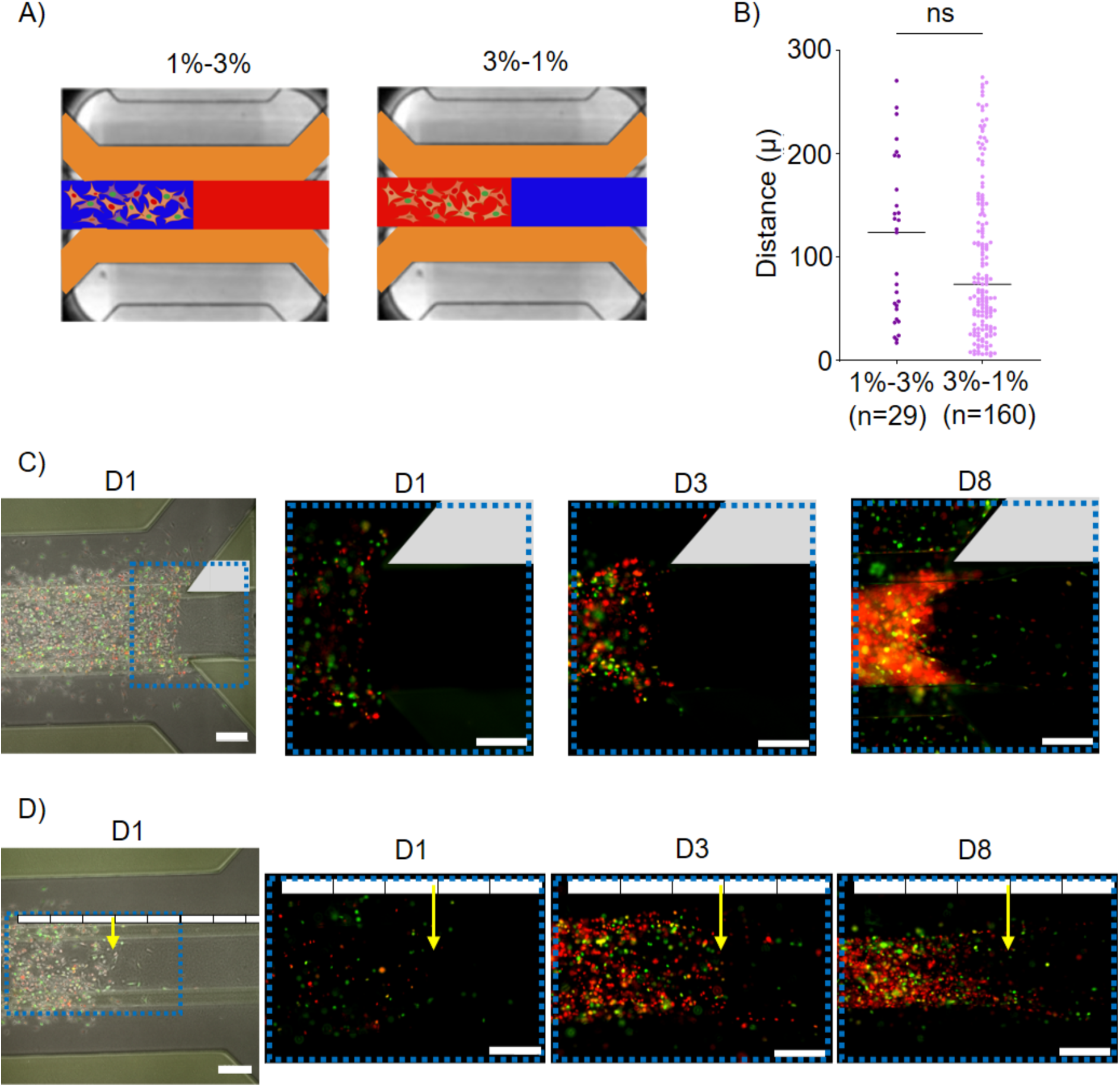
A) Schematic representation of the interphase between 1% (blue) adipose tissue dECM-derived hydrogels embedded with cells and 3% (red) cell-free adipose tissue dECM-derived hydrogels, and vice versa, 3% adipose tissue dECM hydrogels with cells and cell-free 1%, in the central channel. B) Quantification of the invaded distance in the cell-free adipose tissue dECM-derived hydrogel for the MDA-MB-231 cells (each dot represents an invading cell) at day 8 of culture, using Fiji (right). The migration from the 3% into the 1% hydrogel (light purple) and from the 1% into the 3% hydrogel (dark purple) were analyzed using an unpaired Student’s t-test: n.s (p > 0.05). C) Bright field and fluorescence microscope images of MDA-MB-231 cells expressing FUCCI2 (G1: red, G2/M: green) in the area of interest, including regions from the 1% and the 3% adipose tissue dECM-derived hydrogels and their interphase (1%-3%) after 1, 3 and 8 days. The grey trapezoid highlights the structural guide used as a reference for cell migration. The blue dashed line indicates the zoom-out area imaged at higher magnification captured at subsequent timepoints. Experiments were repeated two times per condition. Scale bar: 200 μm. D) Bright field and fluorescence microscope images of MDA-MB-231 cells expressing FUCCI2 (G1: red, G2/M: green) in the area of interest, including regions from the 3% and the 1% adipose tissue dECM-derived hydrogels and their interphase (3%-1%) after 1, 3 and 8 days. The white rectangles, each 200 μm long, mark the distance from the end of the middle channel, used as a reference for cell migration. The yellow arrow marks the interphase between the 3% and 1% hydrogels. The blue dashed line indicates the zoom-out area imaged at higher magnification captured at subsequent timepoints. Experiments were repeated two times per condition. Scale bar: 200 μm.

Our experiments reproduced both, (i) the well-characterized behavior of luminal, poorly aggressive breast cancer, modeled with MCF-7 cells, and (ii) the triple-negative, highly metastatic breast cancer, modeled with MDA-MB-231 cells. MCF-7 cells proliferated within the adipose tissue dECM-derived hydrogels but did not invade the adjacent cell-free gel (Figures 7B and C). Notably, they exhibited a tendency to leave the hydrogel and grow in 2D in the perfusion channels (Figure S5A and B). In contrast, MDA-MB-231 cells both proliferated and invaded the adjoining 1% and 3% acellular hydrogels (Figures 8C and D). Consistent with the proliferation findings described earlier for this cell line, MDA-MB-231 cells that invaded the cell-free gel subsequently proliferated more in the adjoining 1% adipose tissue dECM-derived hydrogel than in the 3% (Figure 8D). Interestingly, there was no statistical difference in the distance invaded in gels without cells at both 1% and 3%.

## 3. Discussion

In this study, we used adipose tissue dECM-derived hydrogels to model the mechanical characteristics and structural protein composition of the breast microenvironment and investigate potential triggers of behavioral changes of breast cancer cells. Our findings reveal that cell proliferation, growth pattern, and invasiveness are highly dependent on both the elastic and viscoelastic properties of the ECM. These properties consistently influenced two well characterized cell lines with distinct behaviors, MCF-7 and MDA-MB-231.

The composition and structure of the ECM mediate mechanotransduction signaling. Adipose tissue derived dECM hydrogels constitute biologically relevant ECM, providing a natural environment for cells while retaining both the native biochemical and the biophysical properties of the source tissue. In particular, these hydrogels allow thermally induced gelation, thereby avoiding the need to expose cells to UV light, which can induce DNA damage even with short exposures. ^[38]^ This is particularly crucial when working with cancer cell lines, as UV light is a carcinogen and DNA damage is a hallmark of cancer. ^[39]^ Additionally, adipose tissue dECM-derived hydrogels can be tailored to meet the specific physicochemical requirements of research, offering advantages over other materials such as collagen hydrogels and Matrigel. Moreover, these hydrogels demonstrate superior long-term stability compared to Matrigel and, being non–tumor-derived, help reduce variability and ethical concerns. ^[13]^ The proteomic analysis revealed that, as expected, most of the identified proteins in the adipose tissue dECM-derived hydrogels were related to collagen, which is a major structural component of the ECM. The protein abundance profile closely resembled that observed in matrisome analysis of human adipose tissue-derived dECMs. ^[40]^ Proteins involved in cell adhesion, such as lectin and laminin, were detected as well. These molecules are essential for integrin function and are known to promote endothelial and epithelial cell adhesion. ^[41]^ Notably, confocal reflectance imaging revealed preservation of the fibrous architecture of the ECM within these adipose tissue-derived dECM hydrogels.

The adipose tissue dECM-derived hydrogels we employed mimic breast tissue stiffness presenting complex moduli *G\** of 60 Pa and 300 Pa for the 1% and the 3% hydrogels, respectively. This corresponds to elastic moduli E of 180 Pa for the 1% hydrogel and 900 Pa for the 3% hydrogel. These values fall within the healthy range for breast tissue, which has storage moduli between 160 Pa to 950 Pa ^[1–3]^ although they can vary depending on the measuring technique and the composition of the tissue (with fat being significantly stiffer than the fibroglandular component). The phase angle tangent takes values above zero, indicating some energy dissipation, as found in actual tumor samples. ^[42]^ The stress relaxation half-times have mean values of 2.3 s and 14.8 s for the 1% and 3% hydrogels, respectively. As a note of caution, even if our stress relaxation times are lower than those reported by others, who consider fast relaxing (*τ*_1/2_) = 30 s ^[43]^, 80 s ^[44]^ or even 3 min, ^[17]^ these numbers are not absolute measures. Beyond the lack of standardization in the protocols for mechanical characterization of tissues and hydrogels, the variation is further compounded by the use of different biomaterials (with widely varying elastic moduli), fabrication techniques, and modes of mechanical characterization (e.g., stress relaxation tests performed in both shear and compression modes), which might explain the discrepancies.

The differences in elastic moduli of the two adipose tissue dECM-derived hydrogels, with 180 Pa for the 1% gel and 900 Pa for the 3% gel, show statistical significance. Nevertheless, they differ by less than 800 Pa, a relatively small variation compared to the multi-kilopascal differences commonly reported in hydrogel studies, with ranges of 1.4-9 kPa, ^[18]^ 3-16 kPa, ^[44]^ 0.4-9.3 kPa. ^[45]^ Yet, despite this modest stiffness range, which falls within the category of ‘soft’ hydrogels in the field, MCF-7 and MDA-MB-231 cells exhibited markedly different behaviors when cultured in 1% versus 3% hydrogels. We hypothesized the involvement of another mechanical property. Giving the evidence suggesting tissue viscoelasticity may correlate with cancer progression, ^[19]^ we investigated the stress relaxation of the adipose tissue dECM-derived hydrogels. Our findings indicate that both MCF-7 and MDA-MB-231 cells exhibit increased proliferation, with MDA-MB-231 cells also presenting enhanced invasiveness towards the softest and fastest relaxing 1% adipose tissue dECM-derived hydrogels. These results are consistent with those reported by Nam et al. ^[44]^ for those same breast cancer cells and by Sinha et al. ^[46]^ for glioblastoma cells. Intriguingly, the growth pattern appears to depend on ECM properties rather than being inherent to the cancer cells themselves, as cells changed their proliferation rate upon crossing the interphase boundary. This change occurred regardless of whether the cells moved from a soft to a stiffer hydrogel or vice versa. Aligned also with the findings of Nam et al. ^[44]^, the fast stress-relaxing 1% adipose tissue dECM-derived hydrogels promoted cell cycle progression in both MCF-7 and MDA-MB-231 cells. In fact, this rapid stress relaxation due to matrix remodeling has been linked to metastasis. ^[47]^ Conversely, MCF-7 cells showed a tendency towards growth arrest in the stiffer and slower relaxing 3% hydrogels, as indicated by a higher G1 to G2/M ratio.

Despite the seemingly high heterogeneity among tumors, several authors have pointed out that tumor tissue exhibits structural patterns corresponding to organ developmental stages. ^[48]^ In both healthy and cancerous tissues, cells do not exist in isolation but rather form fluid cellular aggregates that collectively sense and respond to stimuli. These stimuli are integrated by biomolecular and physical cues from the ECM environment, which regulates collective cell dynamics and tissue patterning. ^[43,49]^ Our results indicate that cells growing in adipose tissue dECM-derived hydrogels tend to form aggregates. This biologically relevant physicochemical environment enabled MCF-7 and MDA-MB-231 cells to recapitulate their characteristic growth patterns, modelling luminal and triple-negative breast cancer, respectively: MCF-7 cells spontaneously formed spheroids from single cells that increased in size over time, whereas MDA-MB-231 cells exhibited an invasive, polarized migration. This is in accordance with previous studies indicating that cell proximity enhances migratory and invasive behavior, ^[50]^ suggesting an advantage to collective invasion. One could argue that cells are migrating on oriented fibers, as observed by other researchers. ^[51]^ However, our confocal reflectance images of the adipose tissue dECM-derived hydrogels after one day in culture do not show any preferential direction of the fibers. Nevertheless, we do not exclude the possibility that leader cells could alter the structure of the fibers over time, thereby enabling directional invasion.

Most research has focused on measuring the proliferation and invasion capacity of cancer cells within hydrogels with a storage modulus ranging from well above 4 kPa up to 20 kPa. ^[13,31]^ However, most cancer cells migrate not across tumors, but through adjacent interstitial tissue, which is much softer. Other groups have shown that within 1.8 kPa hydrogels (interpenetrating alginate networks in this case), which are similar in stiffness to malignant breast tumors, breast cancer cells extend invadopodia to mechanically open channels for migration. ^[52]^ However, increased stiffness decreases hydrogel plasticity and physically restricts invadopodia extension and cell migration. ^[45]^ Although using a different biomaterial and in another stiffness range, we see a similar behavior, with MDA-MB-231 cells migrating and invading more readily toward the softest and fastest-relaxing adipose tissue dECM-derived hydrogels, specifically from 3% to 1%. Noticeably, although fewer cells invaded across the interface from the 1% to the 3% hydrogel than in the reverse condition, those that did invade advanced a similar distance into the cell-free hydrogel in both conditions. This unexpected result may be attributed to the chip’s structure, where nutrients are more abundant in the center (where the middle channel contacts the perfusion channels), causing cells to be reluctant to migrate away from the hydrogel in contact with the medium (toward the hydrogel inlets on the sides), or to the intrinsic migratory capacity of the MDA-MB-231 cells. Invasive MDA-MB-231 cells could migrate through the interface between hydrogels, suggesting that the matrix boundary of the interface did not hinder cell migration.

All things considered, our findings highlight the critical role of the biochemical and biophysical microenvironment in shaping tumor cell phenotypes, which respond to matrix viscoelastic, elastic and density cues, thereby adapting their behavior. Understanding and monitoring these key values could enhance predictive tools and personalized medicine.

## 4. Conclusion

Two distinct breast cancer cell lines mimicking luminal and triple negative breast cancer exhibited strong behavioral differences in adipose tissue-derived dECM hydrogels. Soft and viscoelastic adipose tissue dECM hydrogels favored proliferation, migration and invasion. Furthermore, we could recapitulate distinct collective growth patterns characteristic of these two breast cancer types. This suggests that a combination of various biophysical properties, such as stiffness and stress relaxation, among others, leads to phenotype switching. Our relatively simple and user-friendly model can recapitulate crucial aspects of in vivo behavior under certain conditions.

## 5. Experimental Section/Methods

### 5.1. Adipose tissue decellularized extracellular matrix-derived hydrogel fabrication

The dECM obtained form porcine adipose tissue was manufactured and supplied by Tecnalia, in particular material from batch PPD3-1-22, as previously described. ^[30]^ This dECM was obtained by homogenization and treated with an organic solvent and a detergent (leaving only traces of DNA or lipids). Finally, it was lyophilized and ground into a powder as described before. ^[30]^ to produce dECM-derived hydrogels, 30 mg/ml of the batch PPD3-1-22 dECM were digested with pepsin (107192, Sigma-Aldrich) at 3 mg/ml under acidic conditions (HCl 0.1 N, Honeywell Fluka) for 48 h at room temperature. Finally, the pH was neutralized with NaOH 1 N (Honeywell Fluka) to pH 7.3 on ice and phosphate-buffered saline (PBS) 10x (14200075, Gibco) was added to adjust the salt concentration. The resulting pregel solution at 3% w/v, mixed or not with cells depending on the experiment, was diluted with PBS 1x (10010015, Gibco) to a final concentration of 1% w/v or kept at 3% w/v (referred to as 1% and 3%, respectively, from now on). It was poured into custom-made Teflon rings, 2 mm in height and 8 mm in diameter (for cell culture) or 2 mm in height and 20 mm in diameter (for rheology). It was thermally gelled by keeping it for 50 min in the cell culture incubator at 37°C, 5% CO2 and 95% humidity.

### 5.2. Proteomic analysis of the source dECM powder and dECM-derived hydrogels

The relative quantification of the proteins in the dECM was characterized by label free liquid chromatography-mass spectrometry (LC-MS/MS) at the Proteomic Platform at CIC bioGUNE (Derio, Spain). Triplicates of two types of samples were characterized: the source adipose tissue dECM powder from Tecnalia and the 3% w/v adipose tissue dECM-derived hydrogel. Sodium dodecyl sulfate-polyacrylamide gel electrophoresis (SDS-PAGE) was performed on these samples at three different concentrations to visualize protein sizes (Figure S6). After lyophilization and grinding into a powder of the 3% hydrogel, both samples were resuspended in a buffer containing 7M urea, 2M thiourea, 4% 3-[(3-cholamidopropyl) dimethylammonio]-1-propanesulfonate (CHAPS) and 200 mM dithiotreitol (DTT). For the digestion of the samples, approximately 20 µg were processed following the filter-aided sample preparation (FASP) protocol described previously. ^[53]^ Resulting peptides were desalted and resuspended in 0.1% formic acid using C18 stage tips (Millipore). They were analyzed in a hybrid trapped ion mobility spectrometry – quadrupole time of flight mass spectrometer (timsTOF Pro with PASEF, Bruker Daltonics) coupled online to a Evosep ONE liquid chromatograph (EVOSEP). Approximately 200 ng were loaded for each sample. Samples were resolved using the default 30 samples-per-day (SPD) protocol in the chromatograph and acquired in data-dependent acquisition (DDA) mode in the mass spectrometer using default methods.

Protein identification and quantification were carried out using PEAKS X+ software (Bioinformatic Solutions) with 20 ppm and 0.05 Da tolerances for precursor and fragment masses, respectively. Carbamidomethylation of cysteines was considered a fixed modification, whereas oxidation of methionine and hydroxylation of proline were considered variable modifications. The search considered proteins cleaved with trypsin with a ragged end, and was carried out using Uniprot reference proteome with pig (Sus scrofa) protein entries. Area was used for the relative quantitative analysis. These values were normalized to the total signal of each sample, to be able to express them as %, and compared the groups using a t-test. To avoid redundancy, work was done at the gene level, selecting only a group of values for each gene. Results were represented on a volcano plot generated with VolcanoSer (https://huygens.science.uva.nl/VolcaNoseR/).

A gene ontology (GO) functional analysis was performed using the DAVID program (https://davidbioinformatics.nih.gov/) for those genes identified in at least 2 out of three replicates of the dECM-derived hydrogel samples, of the source dECM powder samples and differentially on the hydrogel sample vs. source dECM powder (>2 fold). A manual curation of the data was conducted, discarding intracellular and membrane proteins (artifacts from cellular debris) and keeping ECM-related proteins.

A sodium dodecyl sulfate-polyacrylamide gel electrophoresis (SDS-PAGE) of the two different sample types was carried out at several concentrations (3.5 µg, 7 µg and 9.3 µg) to visualize differences in the molecular weight of the proteins. The gel was stained with SimplyBlue SafeStain (LC6060, Invitrogen) Coomassie.

[The raw data corresponding to the proteomic analysis in this paper is available free of charge in the folder corresponding to Figure 1 on Zenodo, accesible via 10.5281/zenodo.17610161]

### 5.3. Rheological study of elastic and viscoelastic properties of adipose tissue dECM-derived hydrogels

Measurements to assess the rheological properties of the 1% and 3% adipose tissue dECM-derived hydrogels were performed on the ARES-G2 rheometer (TA Instruments) equipped with a 20 mm diameter parallel plate geometry. Experiments were carried out at a constant temperature of 37°C. Samples were kept hydrated during the measurements. The gap was adjusted each time to the height of the gel to maintain a constant normal force of 1N.

Strain sweep tests were performed to determine the linear viscoelastic region, where both the storage (*G′*) and loss (*G″*) moduli are independent of the applied strain. Based on the results of those tests, frequency sweep tests (from 0.02 to 100 Hz) of the samples were performed at a constant strain of 0.5% (n=6 for 1% hydrogels and n=3 for 3% hydrogels). Complex modulus and phase angle tangent of adipose tissue dECM-derived hydrogels were calculated from the frequency sweep data based on *G\** = √ (G′ 2 + G′′2) and tan(δ)= G′/G′′, respectively. Then, the elastic modulus (*E*) was calculated according to the following equation: *E*= 2(1+ ν) G*, based on rubber’s elastic theory and assuming a Poisson’s ratio close to 0.5. ^[54]^

To monitor the stress-relaxation behavior of the adipose tissue dECM-derived hydrogels, a shear strain of 2% was applied to the hydrogels and then the decrease of the stress over time was recorded (n=5 for 1% hydrogels and n=5 for 3% hydrogels). Data were analyzed using the TRIOS Software (TA Instrument) and the time when the stress is reduced to half, *τ*_1/2_, was calculated.

[Further details and raw data for each replicate of the different assays (frequency sweeps and stress-relaxation tests) are available free of charge in the folder corresponding to Figure 2 and Figure S1 on Zenodo, accesible via 10.5281/zenodo.17610161]

### 5.4. Structural characterization of collagen fibers in the adipose tissue dECM-derived hydrogels

Confocal reflectance images of the fibers in the 1% and 3% adipose tissue dECM-derived hydrogels were acquired with the LSM 880 confocal microscope (Zeiss). The objective used was Plan-Apochromat 40x/1.3 NA (Oil DIC M27). The excitation and emission wavelengths of the argon laser were 633 nm and 620-640 nm, respectively. The Z-stacks were obtained as close as possible from the lower edge of the gel and consisted of 67 images with 0.78 μm spacing (50 μm total Z-stack thickness).

[Raw images for each replicate are available free of charge in the folder corresponding to Figure 1 on Zenodo, accesible via 10.5281/zenodo.17610161]

### 5.5. Breast cancer cell culture

Two human breast cancer cell lines were obtained from the American Type Culture Collection (ATCC): MCF-7, ATCC HTB-22 (Lot. 63288596), and MDA-MB-231, ATCC HTB-26 (Lot. 62657852), which serve as models for hormone-responsive, poorly aggressive breast cancer and highly metastatic triple-negative breast cancer, respectively.

Parental cell lines, as well as a genetically modified version of these cells expressing the fluorescent ubiquitination cell cycle indicator 2 (FUCCI2) cell cycle reporter, ^[37]^ previously generated in our lab ^[55,56]^ were used. Briefly, FUCCI2 vectors mCherry hCdt1(30/120) and mVenus hGeminin(1/100) were obtained from the Riken Brain Science Institute and used to generate lentivirus particles by co-transfecting HEK 293TN cells with packaging and envelope plasmids. MDA-MB-231 and MCF7 cell lines were transfected with mVenus hGeminin(1/110) at multiplicities of infection (MOIs) of 6 and 5, respectively, followed by mCherry hCdt1(30/120) at a MOI of 3. Successful transduction and sorting of stably expressing cell lines were verified using flow cytometry. This modification allows for distinguishing between the G1 phase, with the nucleus in red (mCherry-hCdt1), and the G2/M phase, with the nucleus in green (mVenus-hGeminin), without the need for additional staining.

All cell lines were cultured in dulbecco’s modified eagle’s medium (DMEM) (41966029, Gibco) supplemented with 10% v/v fetal bovine serum (FBS) (A5256701, Gibco) with 1% penicillin/streptomycin solution (15140122, Gibco). The cultures were incubated in standard tissue culture conditions at 37°C with 5% CO2.

### 5.6. Breast cancer cell proliferation

MCF-7 and MDA-MB-231 cells were hydrolyzed from 2D culture and centrifuged. Once the supernatant was completely removed, the cells were embedded in 1% and 3% adipose tissue dECM pre-gels by directly mixing them to a final concentration of 250.000 cells/ml. The hydrogels were allowed to self-assembled for 50 min in the cell culture incubator at 37°C, 5% CO2, and 95% humidity. The cells were grown in the dECM-derived hydrogels for 15 days, with images taken on days 1, 5, 8, 12, and 15 with the AxioObserver 7 inverted epifluorescence microscope (Zeiss). Experiments were repeated three times per time point, hydrogel type and cell line. Breast cancer cell proliferation was assessed by measuring the increase in cell number based on the FUCCI2 fluorescence signal, as explained in section 2.10.1.

[Raw data and images from all replicates for each cell line, time point, and hydrogel condition are available free of charge in the folder corresponding to Figure 3 (MCF-7) and 4 (MDA-MB-231) on Zenodo, accesible via 10.5281/zenodo.17610161]

### 5.7. Breast cancer cell growth pattern: MCF-7 spheroid size and number, and MDA-MB-231 cell growth directionality

MCF-7 and MDA-MB-231 cells displayed distinct and characteristic growth patterns. MCF-7 cells, representative of luminal breast cancer, formed spheroids that increased in number and size over time, whereas MDA-MB-231 cells, which model triple-negative breast cancer, showed an invasive growth pattern. Therefore, they were studied separately.

MCF-7 cells embedded in 1% and 3% adipose tissue dECM-derived hydrogels at a concentration of 250.000 cells/ml were grown for 15 days. Images were taken on days 1, 8, and 15 with the AxioObserver 7 inverted epifluorescence microscope (Zeiss). Experiments were repeated four and three times for 1% and 3% dECM-derived hydrogels, respectively. MCF-7 cells grew forming spheroids, as detected by the FUCCI2 fluorescent signal. Spheroid size and number were analyzed as explained in section 2.10.2.

MDA-MB-231 parental cells embedded in 1% and 3% adipose tissue dECM-derived hydrogels at a concentration of 250.000 cells/ml were cultured for 15 days. On days 1, 8 and 15 cells were stained with DAPI/Phalloidin to visualize the nucleus and architecture of the cytoskeleton, respectively. MDA-MB-231 cells exhibited an invasive growth pattern, with invadopodia oriented in a defined direction, as evidenced by phalloidin staining of the cytoskeleton. For DAPI/Phalloidin staining, cells were fixed with 4% paraformaldehyde in PBS, permeabilized with 0.1% Triton X-100 in PBS and stained with a solution containing DAPI (4’,6-diamidino-2-phenylindole) (MBD0015, Merck) and Phalloidin conjugate (Alexa Fluor 488, #8878, Cell Signaling Technologies) at 1:1000 and 1:200, respectively, for 30 minutes in a rocker at room temperature to facilitate dye diffusion and covered with aluminum foil. Following PBS washing, the hydrogels were imaged at the LSM 900 confocal microscope (Zeiss). Experiments were repeated three times per time point and hydrogel type. Microscope images were analyzed as explained in section 2.10.2.

[Raw data for measurements and raw images from all replicates for both cell lines, time point, and hydrogel conditions are available free of charge in the folders corresponding to Figure 5 and Figure S2 (MCF-7), and Figure 6 and Figure S3 (MDA-MB-231) on Zenodo, accesible via 10.5281/zenodo.17610161]

### 5.8. Breast cancer cell migration and invasion capacity

The migration and invasion capacity of breast cancer cells was evaluated using the commercially available, perfusable microfluidic platform from Mimetas. The OrganoPlate® 3-lane (4004-400-B) chip consists of a central channel, used as a hydrogel inlet in our experiments, and two perfusion channels located at the top and bottom (Figure S4A). MCF-7 or MDA-MB-231 cells were embedded in 1% and 3% pre-gels by directly mixing them to a final concentration of 12.000 cells/µl (recommended by Mimetas). This is a considerably high cell concentration, but it ensures a visualization of the discrete invasion process. Then, while keeping the chip on ice to avoid self-assembling and gelification of the hydrogel, 5 µl of cell-laden pregel were introduced through the right inlet of the central channel, and additional 5 µl of either cell-laden or cell-free pregel, depending on the experiment (see below), were introduced through the left inlet of the central channel. This volume is approximately double the manufacturer’s recommendation, owing to the high viscosity of the pre-gel and the need to ensure complete filling of the central channel.

Four groups were generated according to the adipose tissue dECM-derived hydrogel filling the central channel: (i) cell-laden 1% hydrogel filling the entire central channel; (ii) cell-laden 3% hydrogel filling the entire central channel; (iii) cell-laden 1% hydrogel filling half of the channel and in contact with cell-free 3% gel on the other half; (iv) cell-laden 3% hydrogel filling half of the channel and in contact with cell-free 1% gel on the other half (Figure S4B). The high viscosity of the 3% pre-gel resulted in the interphase between gels being positioned closer to the sides, rather than centrally.

Thermal gelation was induced for 50 min at 37°C to form the hydrogel within the central channel. Then, DMEM supplemented with 10% FBS and with 1% penicillin/streptomycin was added to the top and bottom channels (Figure S4A). Perfusion was initiated by placing the dish on OrganoFlow rocker (Mimetas B.V. #MI-OFPR-L) inside the incubator, tilted at 7-degree angle, which moves side to side every eight minutes. This causes the culture medium to flow back and forth, resulting in gravity-driven perfusion. Cell-laden hydrogels were maintained in culture for 15 days at 37°C with 5% CO2. Images of the cells were acquired after 1, 5, 8 and 15 days with the AxioObserver 7 inverted epifluorescence microscope (Zeiss). Experiments were repeated twice per condition and cell line.

[Raw data and images from all replicates for each cell line, time point, and hydrogel condition are available free of charge in the folders corresponding to Figure 7 (MCF-7), Figure 8 (MDA-MB-231), and Figure S4 and S5, on Zenodo, accesible via 10.5281/zenodo.17610161]

### 5.9. Image acquisition

#### 5.9.1. Fluorescence microscopy of FUCCI2 breast cancer cells within adipose tissue dECM-derived hydrogels

Proliferation and invasion images (sections 2.6, 2.7 and 2.8) were acquired with the AxioObserver 7 inverted epifluorescence microscope (Zeiss). The objectives used were EC Plan-Neofluar 5x/0.16 NA Ph1 M27 or 10x/0.30 NA Ph1. For the 5x magnification, the fluorescence channels mCherry and mVenus were recorded at 60% and 70% LED intensity at 587 nm and 524 nm excitation wavelength, respectively. For the 10x magnification, mCherry and mVenus were recorded at 25% and 70% LED intensity at those same excitation wavelengths. 45 Texas Red Reflector (540/80 excitation and 593/668 emission) was used to image mCherry, and 46 HE YPF Reflector (488/512 excitation and 520/50 emission) was used to image mVenus. Exposure time was 350 ms for mCherry and 340 ms for mVenus. Images of the cells were acquired at day 1, 5, 8 and 15. For proliferation and growth pattern analysis Z-stacks consisting of 10 images with 10 μm spacing (100 μm thickness) were acquired.

#### 5.9.2. Confocal microscopy of MDA-MB-231 cells growing within adipose tissue dECM-derived hydrogels

Cell growth pattern images (section 2.7) were acquired using the Zeiss LSM 900 confocal microscope. The objectives used were Plan-Apochromat 10x/0.45 NA and Plan-Apochromat 63x/1.4 NA (Oil DIC M27). The 405 nm (Blue or DAPI) and 488 nm (green or AF488) lasers were used at 2% power. Emission wavelengths were 465 nm and 517 nm, respectively. The detector was the Multi Alkali-PTM. Z-stacks consisting of 20 images with 5 μm spacing (100 μm thickness) were acquired.

### 5.10. Image analysis

#### 5.10.1. Breast cancer cell proliferation quantification

The increase in cell number was assessed using the FUCCI2 fluorescence signal, based on three orthogonal projections taken from each of three biological replicates per time point and cell line. Data were analyzed using Dragonfly software, Version 2022.1 for Windows (Comet Technologies Canada Inc., Montreal, Canada; software available at https://www.theobjects.com/dragonfly). This software quantifies cell number in two main steps: segmentation and counting. During the first step, a semantic segmentation artificial intelligence model with a U-Net architecture is trained to distinguish between cells and the background using a minimum of 150 cells from various regions of the image, displaying a variety of fluorescent intensities. During the counting step, the model is applied to a single microscope image, sequentially to each fluorescent channel, mCherry and mVenus, corresponding to that image. Cells at the S phase have signal in both mCherry and mVenus channels. To avoid counting these cells twice, they are subtracted from the red count. In order not to consider small particles as cells, spots smaller than 5 µm were removed. Examples of orthogonal projections of microscope images at days 8 and 15 of MCF-7 and MDA-MB-231 cells growing in 1% and 3% hydrogels and their respective Dragonfly masks are shown in Figure S7 and S8.

[The Drangonfly model is available free of charge on Zenodo, accesible via 10.5281/zenodo.17610161]

#### 5.10.2. Breast cancer cell growth pattern analysis

MCF-7 spheroids size and number were analyzed using Dragonfly software, which can quantify clusters (cells that contact each other) and their size in two main steps: segmentation and measuring. During the segmentation step, a mask or region of interest (ROI) is created covering different spheroids but not the background. The artificial intelligence model was refined and optimized by adjusting the ROI over at least five iterations. We analyzed maximum projection images of 100 μm-thick z-stacks from four and three replicates per time point for 1% and 3% adipose tissue dECM-derived hydrogels, respectively. For measuring, we select the 2D equivalent diameter on the scalar generator. In order not to consider small particles or single cells, only clusters bigger than 30 µm were counted (given that the average epithelial cell is 15 µm long, a spheroid must consist of at least two cells). Figure S2 shows orthogonal projection images of MCF-7 cells growing in 1% and 3% hydrogels at day 15 and their respective Dragonfly masks.

Single slices from six different replicates per time point of invasive MDA-MB-231 cells stained with DAPI/Phalloidin were analyzed using Dragonfly software, which can quantify the predominant direction in which cells are growing (directionality) in two main steps: segmentation and measuring. During the segmentation step an artificial intelligence model is trained to distinguish between two regions of interest (ROIs), that is, cells and background. For measuring, we select the 2D orientation on the scalar generator tool, which calculates the principal axes of each ROI, corresponding to the longest axis of the object, called the inertia tensor analysis. Afterwards, it calculates the angle between the X-axis and the major axis of the ROI (theta angle). Figure S3A, B and C show, respectively, confocal microscope images of MDA-MB-231 cell growing in 1% and 3% adipose tissue dECM-derived hydrogels at day 8 and 15, and the masks identifying groups of connected cells with a distinct growth direction at day 15. The predominant growth direction of cells in both 1% and 3% adipose tissue dECM-derived hydrogels was measured, with the predominant growth direction in each 3% hydrogel replicate normalized to 0 degrees to enable combining all replicates.

[Both Drangonfly models and the sessions corresponding to spheroid size measurements (MCF-7) and directionality assessments (MDA-MB-231), as shown in Figure S2 and S3 respectively, are available free of charge on Zenodo, accesible via 10.5281/zenodo.17610161]

### 5.11. Statistical analysis

Bar graphs display the mean value ± standard deviation, while dot graphs show the median. Statistical analysis of normality and equal variances was carried out using the Kolmogorov-Smirnov test. Data fitting a normal distribution were analyzed using an unpaired Student’s t-test for single comparisons of means. Otherwise, the Mann-Whitney U test was employed for non-normal data or samples with unequal variances. Results were plotted using GraphPad Prism 8.3.0 software (GraphPad, San Diego, CA). n.s (p > 0.05), * (p< 0.05), ** (p < 0.01), *** (p< 0.001) and **** (p< 0.0001).

## Acknowledgements

The authors express their gratitude to all members of the Cipitria group, with special thanks to Unai Heras for explaining rheological concepts and for his invaluable introduction to Dragonfly and artificial intelligence models. We also thank the Microscopy Facility at the CIC biomaGUNE, particularly Irantzu Llarena’s support with confocal reflectance experiments. This work was funded by the Gipuzkoa Fellow grant (NSM). A. Cipitria would like to acknowledge funding from IKERBASQUE Basque Foundation for Science, from Fundación Científica Asociación Española Contra el Cáncer (grant LABAE223466CIPI), from the Spanish Ministry of Science and Innovation (MCIN/AEI/10.13039/501100011033/FEDER UE, through grant PID2021-123013OB-I00), and from the European Research Council Consolidator Grant (DORMATRIX, 101123883).

## Data Availability Statement

The data supporting this article have been included as part of the Supporting Information. Besides, other data, including raw proteomic data, microscope images of all the replicates, raw and processed, as well as Dragonfly algorithms are available at Zenodo at [10.5281/zenodo.17610161].

