## Supplementary information for "Stiffness and Viscoelasticity of Adipose Tissue Decellularized Extracellular Matrix Hydrogels Influence Proliferation, Growth Pattern, Migration and Invasion of Breast Cancer Cells"

Supplementary material

- Supplementary Figure 1: Rheology (frequency sweep and stress relaxation) tests and confocal reflectance images.
- Supplementary Figure 2: Orthogonal projections of microscope images of MCF-7 cells at day 15 and their corresponding Dragonfly masks.
- Supplementary Figure 3: Confocal microscope images of MDA-MB-231 cells at day 8 and 15, and their corresponding Dragonfly masks.
- Supplementary Figure 4: Schematic representation of the different hydrogel combinations and their corresponding bright-field and fluorescent microscope images within the OrganoPlate® 3-lane chip.
- Supplementary Figure 5: Schematic representation and fluorescent microscope images of MCF-7 and MDA-MB-231 cells expressing FUCCI2 growing within 1% hydrogels and 3% hydrogels after 1 and 8 days in culture within the OrganoPlate® 3-lane chip.
- Supplementary Figure 6: SDS-PAGE of original dECM material and dECM-derived hydrogel.
- Supplementary Figure 7: Orthogonal projections of microscope images of MCF-7 cells at day 8 and 15 and their corresponding Dragonfly masks.
- Supplementary Figure 8: Orthogonal projections of microscope images of MDA-MB-231 cells at day 8 and 15 and their corresponding Dragonfly masks.

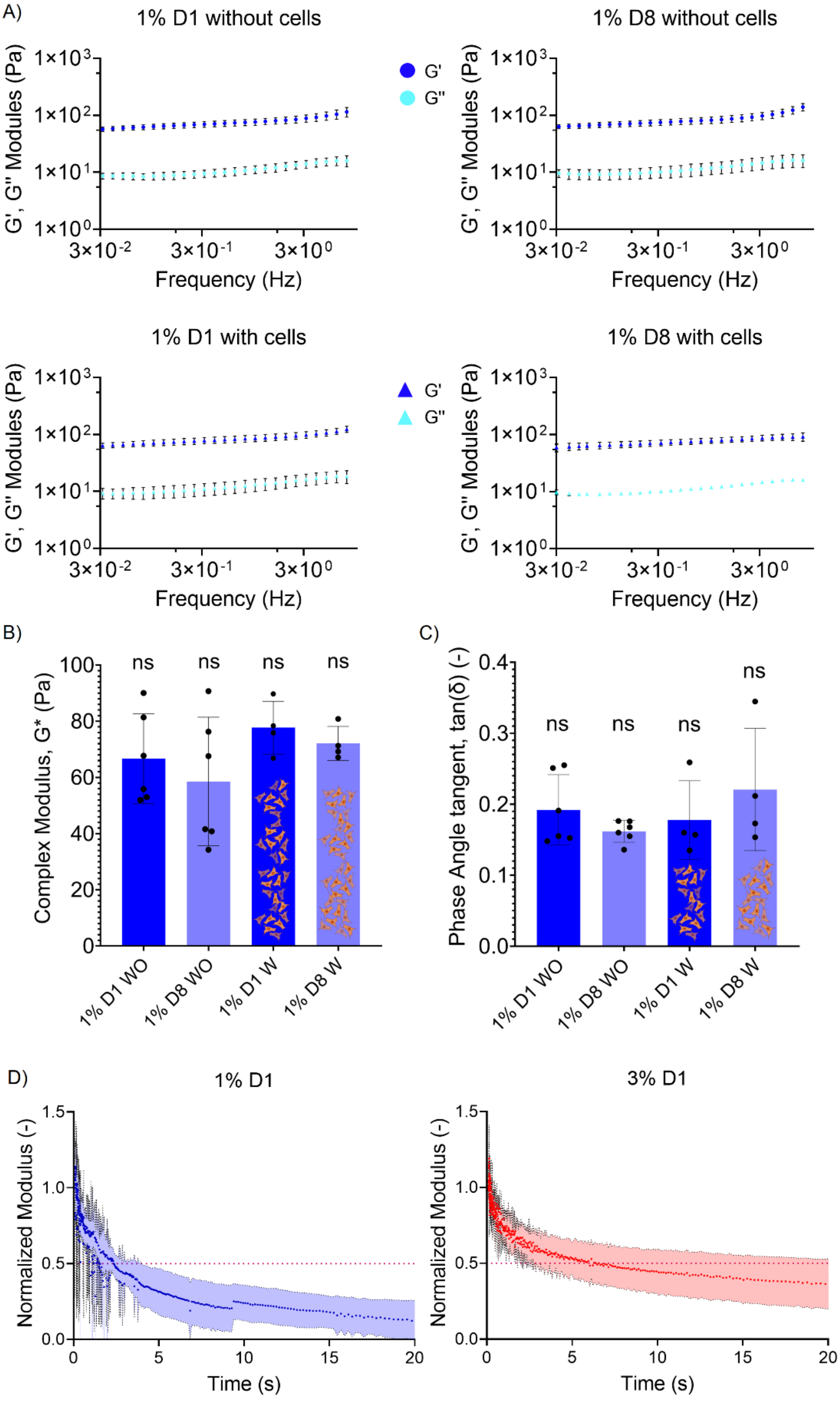


**Supplementary Figure 1**. A) Frequency sweep tests (from 0.03 to 3 Hz) of the 1% adipose tissue dECM-derived hydrogels with (blue triangles) and without (blue circles) cells performed at constant strain (0.5%) at day 1 and 8 days. Experiments were repeated four times for hydrogels with cells and six times for hydrogels without cells. Storage (G′) and loss (G″) moduli are represented as colored and fainter triangles or circles, respectively. B) Complex modulus (G*) and C) phase angle tangent (tan(δ)) of 1% adipose tissue dECM-derived hydrogels with and without cells embedded at day 1 and 8. Data were calculated from the frequency sweep tests in A. D) Normalized individual stress relaxation tests (at 2% shear strain) of the 1% and 3% adipose tissue dECM-derived hydrogels 1 day. n=5 for 1% hydrogels and n=5 for 3%

hydrogels. The dashed lines indicate the time (s) where the modulus is reduced to half, that is, the stress relaxation half-time ($\tau_{1/2}$) of each adipose tissue dECM-derived hydrogel.

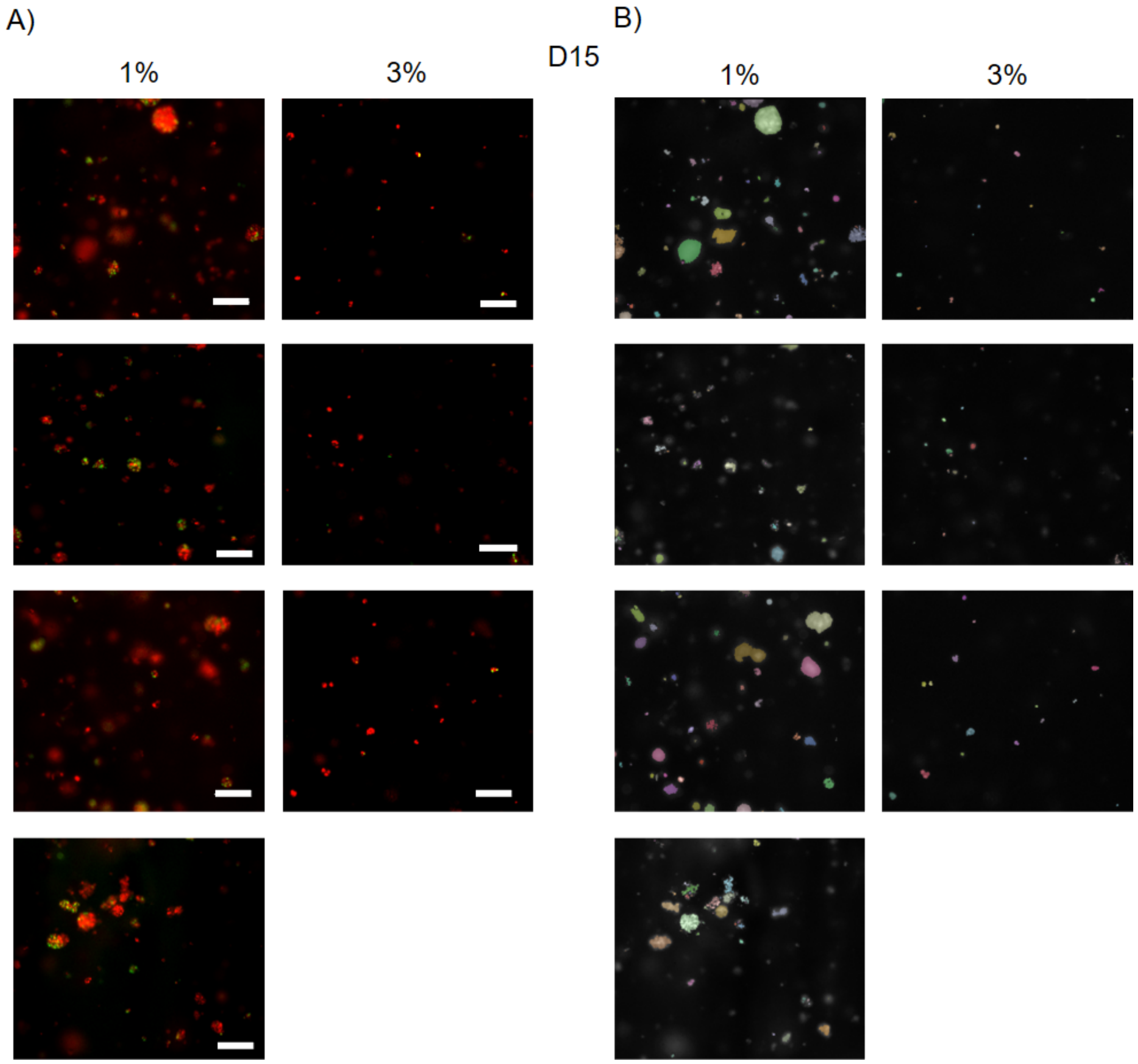


**Supplementary Figure 2**. A) Orthogonal projections of microscope images of MCF-7 cells expressing FUCCI2 (G1: red, G2/M: green) growing within 1% and 3% adipose tissue dECM-derived hydrogels at day 15 shown side by side for easier visual comparison. Experiments were repeated four and three times for 1% and 3% hydrogels, respectively. Scale bar: 100 µm. B) Dragonfly masks corresponding to the images in A) in the same order as the original microscope images.

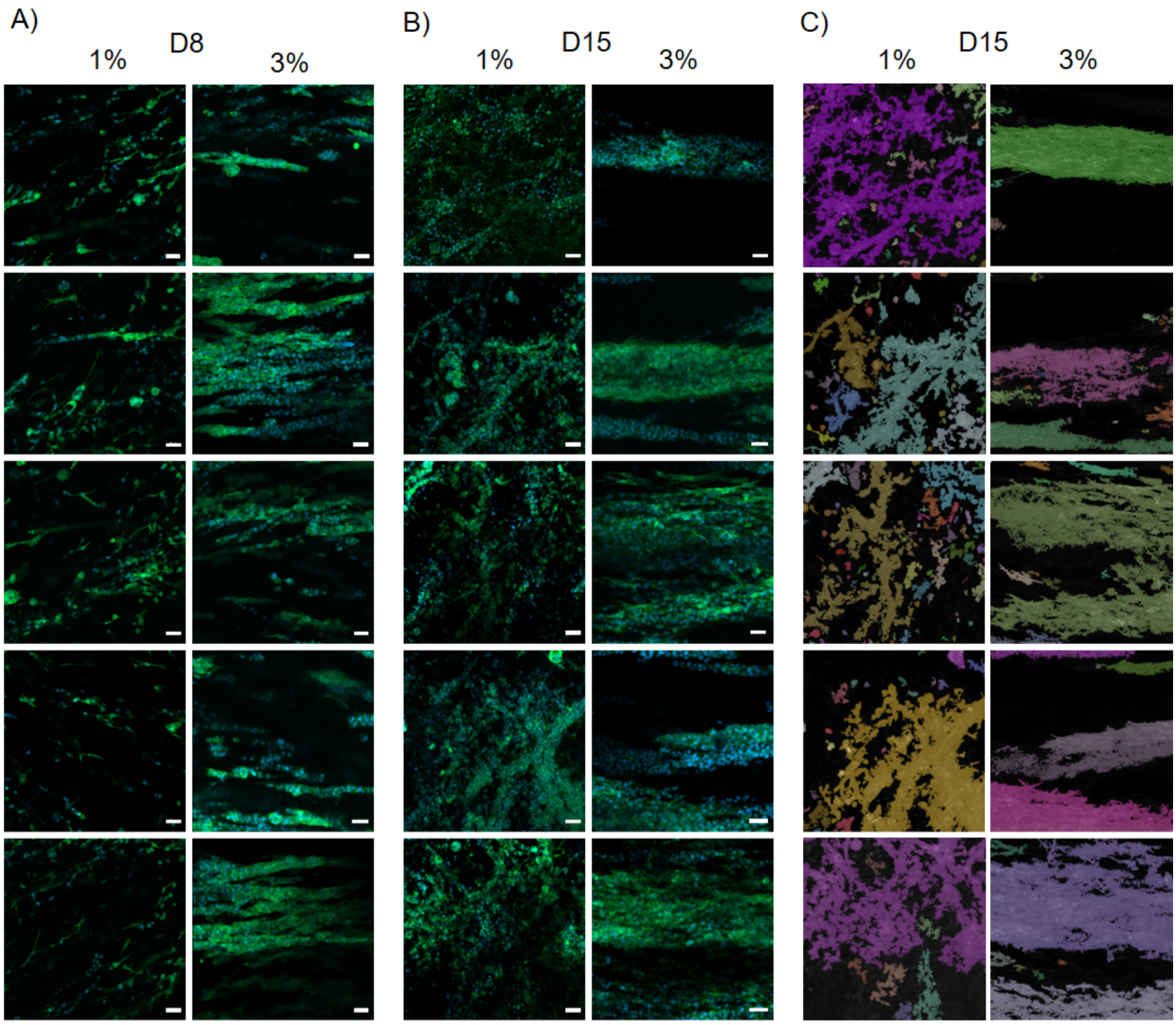


**Supplementary Figure 3**. Confocal microscope images of MDA-MB-231 cells stained with DAPI (blue) and phalloidin (green), growing within 1% and 3% adipose tissue dECM-derived hydrogels after A) 8 days and B) 15 days in culture, shown side by side for easier visual comparison. Experiments were repeated six times for each condition. Scale bar: 50 µm. C) Masks segmented by Dragonfly identifying groups of connected cells (different colors) with a principal direction in 1% and 3% adipose tissue dECM-derived hydrogels at day 15, in the same order as their corresponding original confocal microscope images in B).

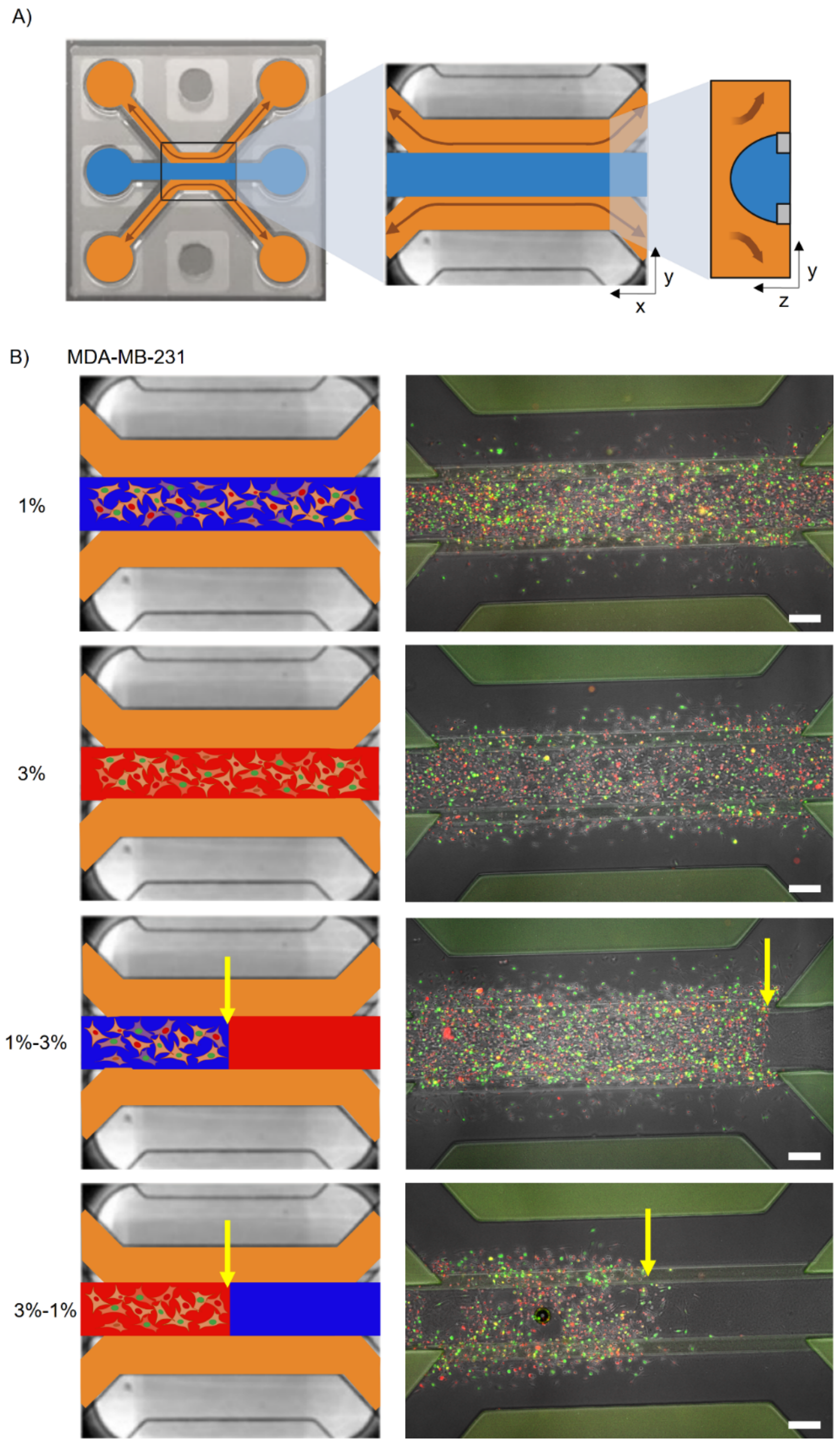


**Supplementary Figure 4**. A) Schematic representation of an OrganoPlate® 3-lane chip. The gel inlet (middle channel) is depicted in blue, and the perfusion channels (top and bottom channels) are shown in orange. The arrows indicate the directions of fluid flow. Views along the X-Y and Y-Z planes are provided. B) Schematic representation of the different hydrogel combinations and their corresponding bright-field and fluorescent microscope images at day 1 of MDA-MB-231 cells expressing FUCCI2 (G1: red, G2/M: green) growing in the following conditions: (i) 1% (blue) adipose tissue dECM-derived hydrogels; (ii) 3% (red) adipose tissue dECM-derived hydrogels; (iii) 1% (blue) adipose tissue dECM-derived hydrogels embedded with cells in contact with 3% (red) cell-free adipose tissue dECM-derived hydrogels; and (iv) 3% (red) adipose tissue dECM-derived hydrogels embedded with cells in contact with 1% (blue) cell-free adipose tissue dECM-derived hydrogels. The yellow arrow indicates the interphase between different hydrogels. Experiments were repeated twice per condition. Scale bar: 200 μm.

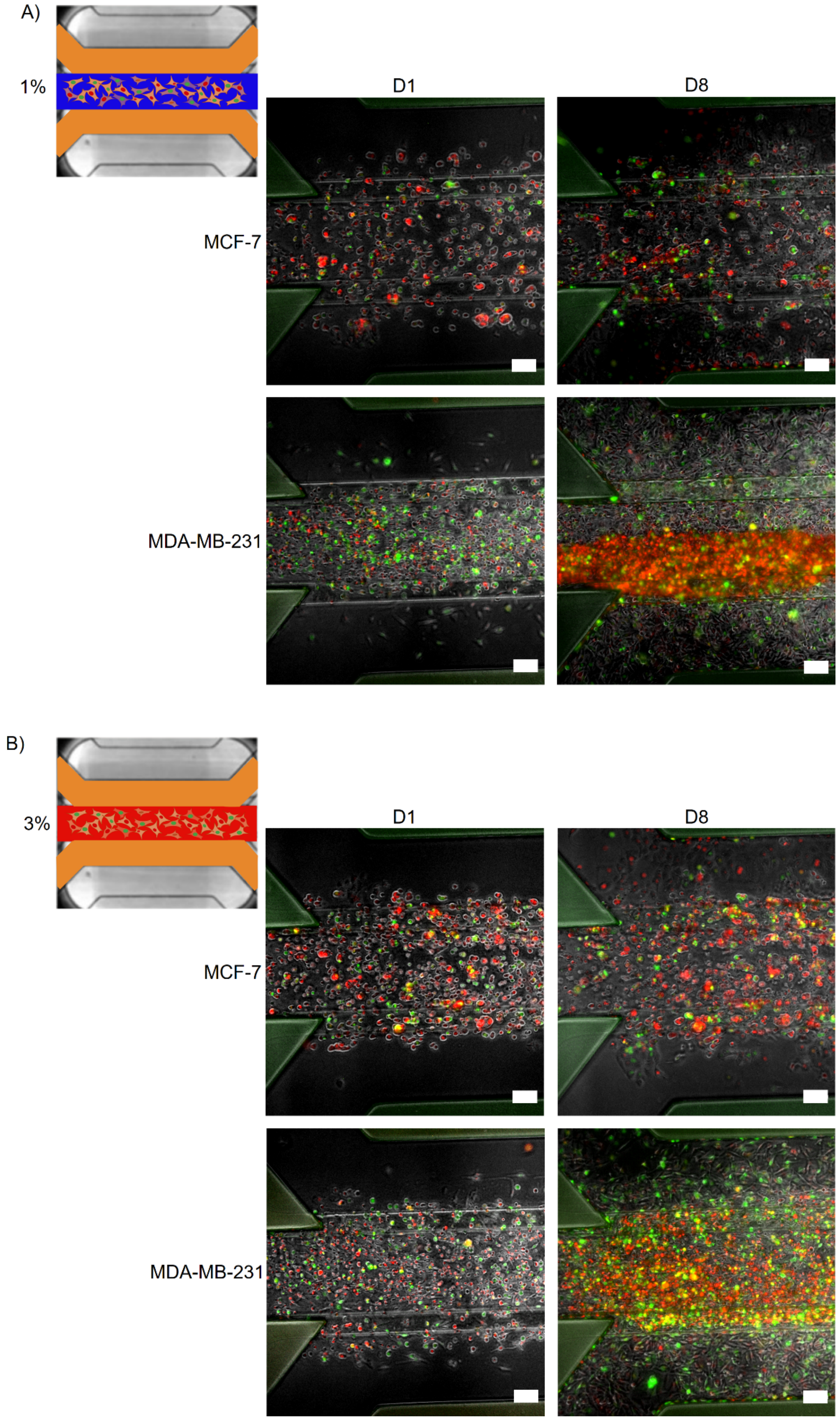


**Supplementary Figure 5**. Schematic representations and fluorescent microscope images of MCF-7 and MDA-MB-231 cells expressing FUCCI2 growing within A) 1% hydrogels and B) 3% hydrogels after 1 and 8 days in culture. Experiments were repeated twice per time point, hydrogel type and cell line. Scale bar: 100 µm.

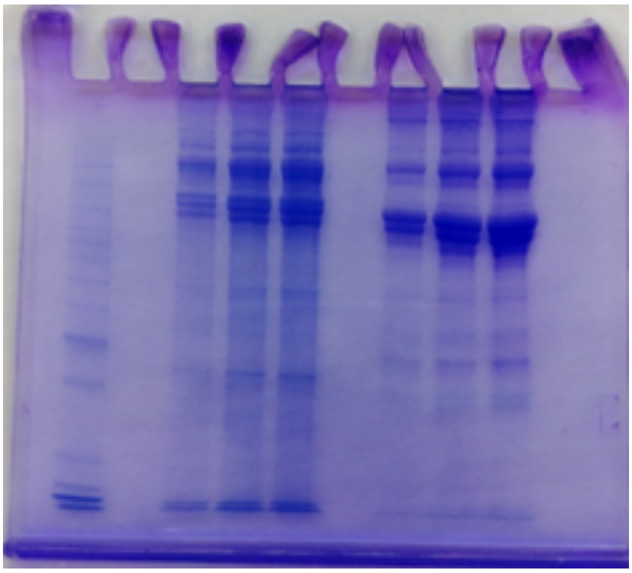

**Supplementary Figure 6**. SDS-PAGE stained with SimplyBlue SafeStain Coomassie. Three concentrations (3,5 µg, 7 µg and 9,3 µg) of each sample type, source adipose tissue dECM powder (1, 2, 3) and 3% adipose tissue dECM-derived hydrogel (4, 5, 6), were analyzed.

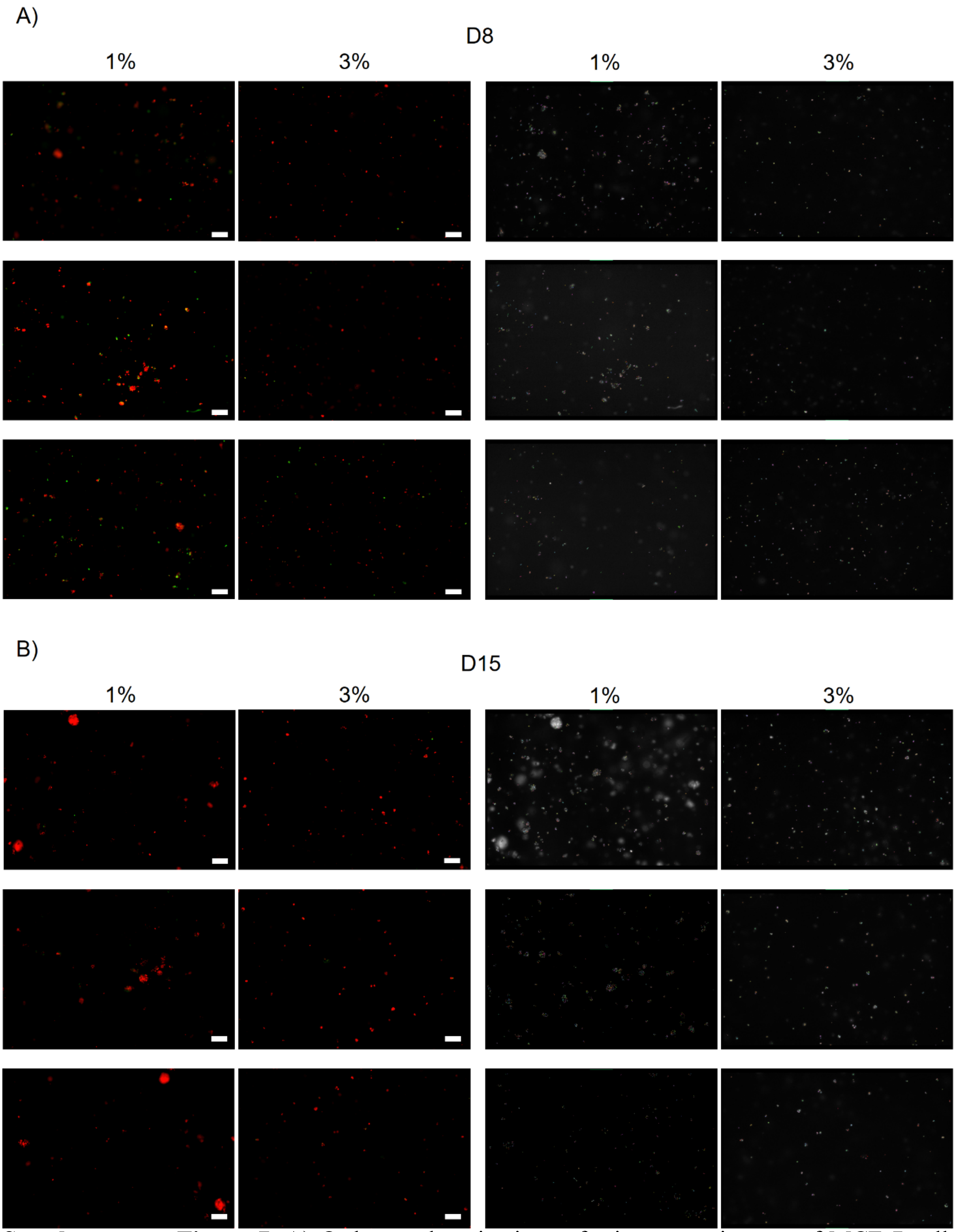


**Supplementary Figure 7**. A) Orthogonal projections of microscope images of MCF-7 cells expressing FUCCI2 (G1: red, G2/M: green) growing within 1% and 3% adipose tissue dECM-derived hydrogels at day 8 shown side by side for easier visual comparison. B) Dragonfly masks corresponding to the images in A) in the same order as the original microscope images. C) Orthogonal projections of microscope images of MCF-7 cells expressing FUCCI2 growing within 1% and 3% adipose tissue dECM-derived hydrogels at day 15 shown side by side for easier visual comparison. D) Dragonfly masks corresponding to the images in C) in the same order as the original microscope images. Experiments were repeated three times for each condition. Scale bar: 200 µm.

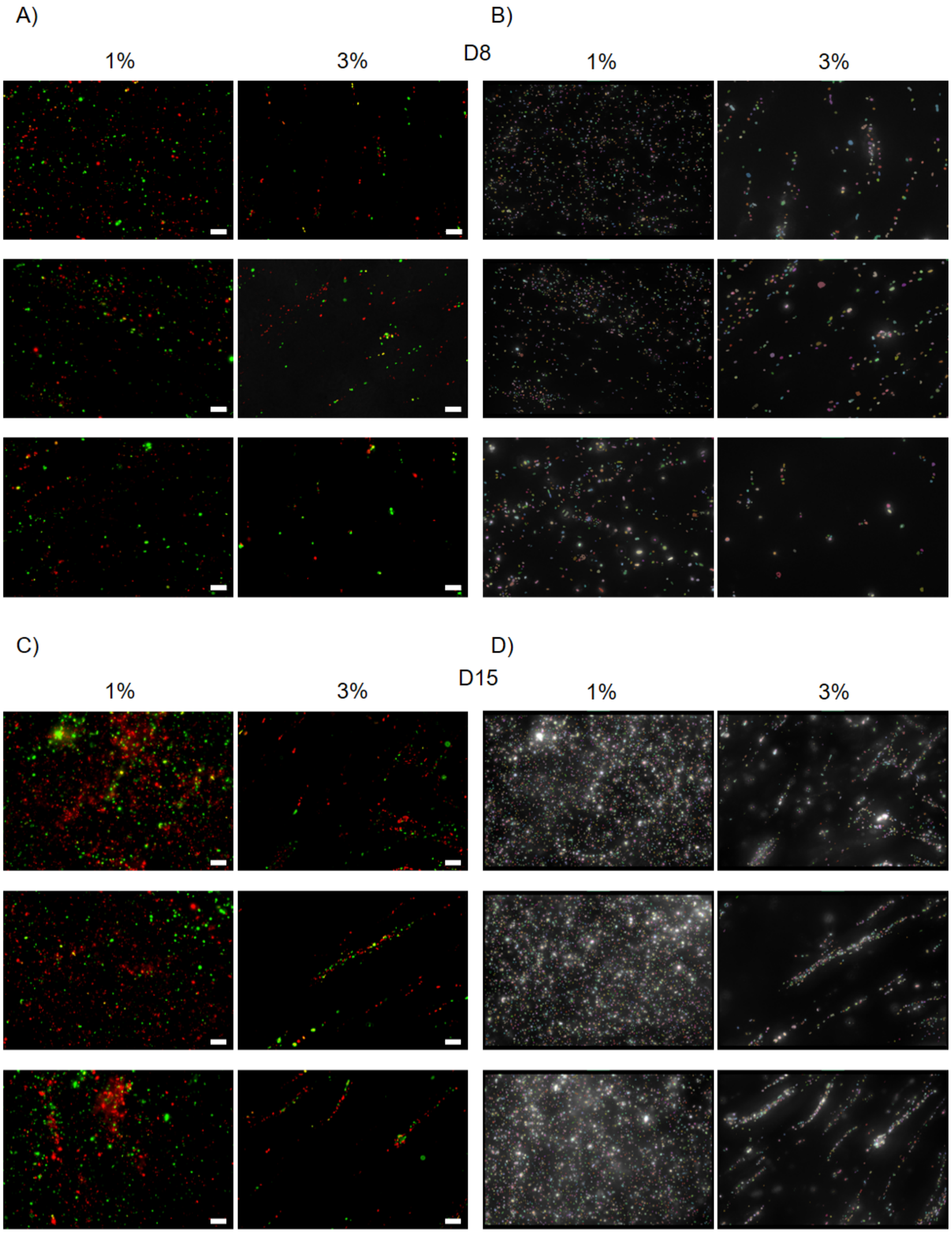


**Supplementary Figure 8**. A) Orthogonal projections of microscope images of MDA-MB-231 cells expressing FUCCI2 (G1: red, G2/M: green) growing within 1% and 3% adipose tissue dECM-derived hydrogels at day 8 shown side by side for easier visual comparison. B) Dragonfly masks corresponding to the images in A) in the same order as the original microscope images. C) Orthogonal projections of microscope images of MDA-MB-231 cells expressing FUCCI2 growing within 1% and 3% adipose tissue dECM-derived hydrogels at day 15 shown side by side for easier visual comparison. D) Dragonfly masks corresponding to the images in C) in the same order as the original microscope images. Experiments were repeated three times for each condition. Scale bar: 100 μm.
